# Viruses help shape microbiome response to polyphenol rewiring of methane-suppressed peat microcosms

**DOI:** 10.64898/2025.12.19.695563

**Authors:** James Riddell, Rokaiya Nurani Shatadru, Garrett J Smith, Bridget B McGivern, Jared B Ellenbogen, Sophie K Jurgensen, Ami Fofana, Malak M Tfaily, Kelly C Wrighton, Matthew B Sullivan

## Abstract

Human activities are accelerating permafrost thaw and subsequent methane emissions from increased microbial activity, prompting microbiome engineering efforts as an emissions mitigation strategy. We recently demonstrated that catechin amendment could drastically reduce methane emissions (>80%) in peat microcosms by enriching catechin-degrading prokaryotes that outcompeted methanogens for hydrogen. However, viral contributions to such microbiome-level responses remain unexplored and we hypothesized that viral dynamics could help shape the microbiome response as nutrient amendments may alter cellular physiology in ways that could induce lytic viral activity. Here, we performed virus ecogenomics analyses of the previously-studied time-resolved multi-omics data collected from catechin-amended peat microcosms. We conservatively identified 900 putatively lytic viral operational taxonomic units (vOTUs), with 41% predicted to infect active host genomes including the most transcriptionally active vOTUs predicted to infect key catechin-degrading genera (*Clostridium* and undescribed *Bacillota* JAGFXR01). Notably, a single JAGFXR01-targeting vOTU dominating the viral response (>40% of community viral transcription; 20-156-fold more abundant than its host), which we interpreted as induction resulting in intense lytic activity that could release catechin-degradation intermediates to other community members. Consistent with this, gene expression analysis revealed elevated catechin-intermediate degradation and hydrogenase signals in 34 additional polyphenol-degrading metagenome-assembled genomes. These findings support a model consistent with a viral shunt-like process that extends our previous prokaryote-centric model: viral lysis of fast-growing catechin degraders redistributes phenolic intermediates to diverse phenol-degrading taxa that sustain methane suppression via hydrogen consumption. Beyond carbon-cycling importance in this system, elucidating unintended virus-mediated responses to nutrient and prebiotic interventions will enable more predictable and effective microbiome engineering strategies across soil, ocean, and human ecosystems.

## Introduction

Microbes form partnerships that are central to nearly all life on Earth, yet anthropogenic climate change is fundamentally altering how these communities function [1]. Warning signs are emerging across the planet: climate-intensified extreme weather events are shifting microbial disease ranges [2], altering coral-microbe symbioses [3], and destabilizing plant-microbe interactions [4–6]. In some systems, climate change may trigger microbial feedback loops that accelerate warming itself. Arctic permafrost thaw exemplifies this risk—as temperatures rise, microbial decomposition of millennia-old soil organic matter releases substantial carbon to the atmosphere [7–9]. At Stordalen Mire (Abisko, Sweden), a long-term, extensively studied subarctic peatland, permafrost thaw over the past 50 years has doubled the extent of fully-thawed fens [10], which emit 4.5 times more methane than intact palsas and partially frozen bogs, driven by increased microbial methanogenesis [11, 12]. As the climate crisis intensifies and microbiome engineering emerges as a potential mitigation strategy [13, 14], identifying interventions to suppress methane production without disrupting fundamental soil microbiome functions has become an urgent priority.

Towards this, recent work at Stordalen Mire has demonstrated catechin, a common polyphenol in Mire flora [15, 16], can decrease methane emissions by >80% in thawed fen peat microcosms [17] (**Figure 1A**). Metagenomics, metatranscriptomics, and metabolomics data collected from these microcosms revealed catechin amendments stimulated community-wide metabolic rewiring where dominant heterotrophic carbohydrate degraders declined, and catechin-degrading taxa, including undescribed *Bacillota* genus JAGFXR01 and *Clostridium* spp., became central to carbon decomposition. Because these newly abundant and active catechin degraders competitively consumed hydrogen and secreted acidic by-products, they collectively suppressed hydrogenotrophic methanogenesis, which is the dominant methanogenesis pathway in these fens [17].

**Figure 1.**
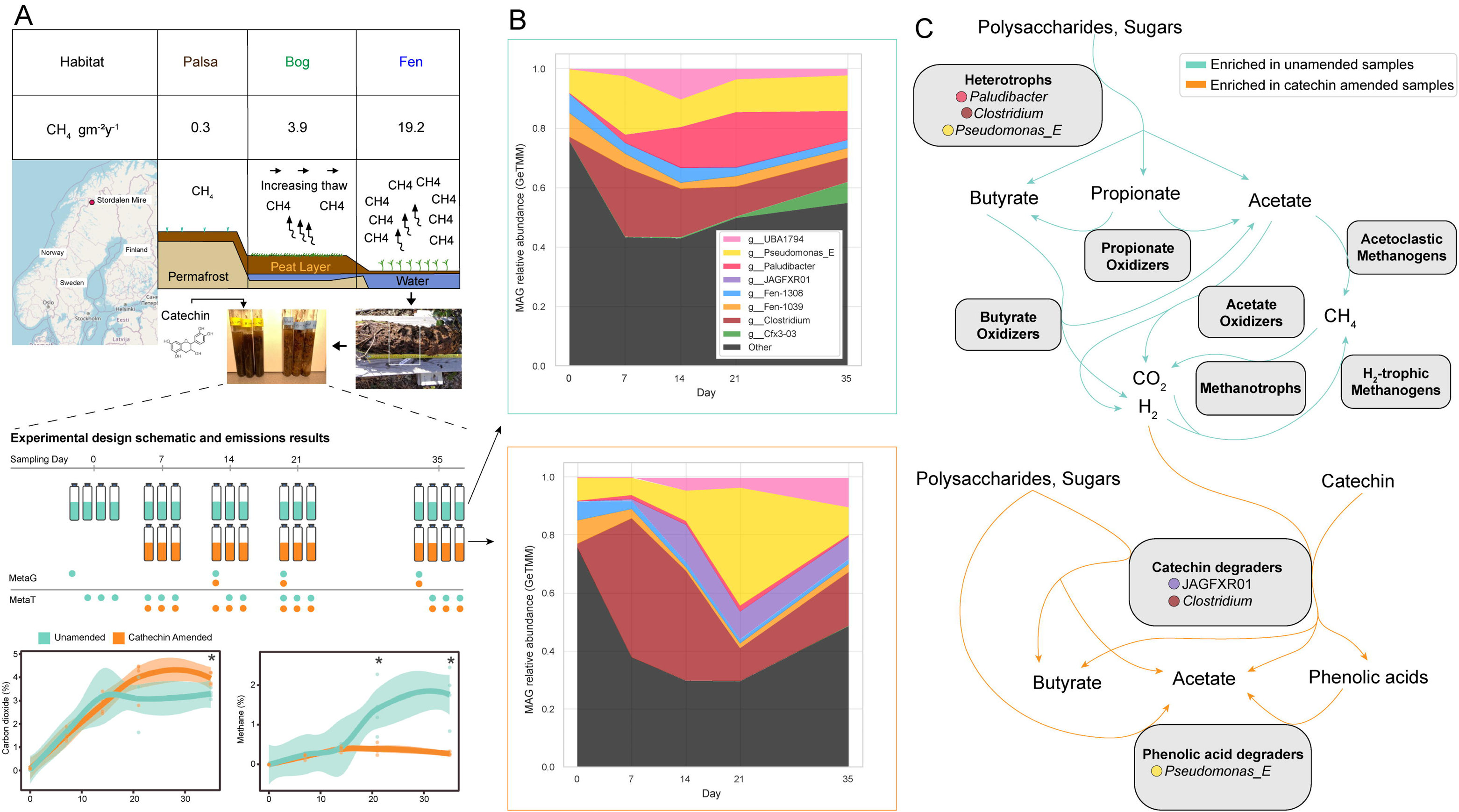
Experimental design overview and summary of prokaryotic community dynamics in response to catechin amendment. **(A)** Peat cores were taken from a fen in Stordalen Mire, Sweden, in July 2016. Microcosms were constructed by placing ∼6g of soil from 9-19cm below the surface (black/white box) into glass Balch tubes. Catechin was added to half of the samples to a final concentration of 4.4 mM, and microcosms were destructively sampled at five timepoints. The sequencing done for each microcosm is specified in the experimental design schematic. Methane flux values derived from [11]. Experimental design schematic and gas measurements adapted [17] under the Creative Commons Attribution license (CC BY). Peat core photo courtesy of Nicole Raab/EMERGE field team (used with permission). Map data © OpenStreetMap contributors (openstreetmap.org/copyright), available under the Open Database License (ODbL). **(B)** MAG community relative activity was visualized by taking the average GeTMM value of each MAG in each set of sample replicates and summing for all MAGs in the same genus. If a genus-level assignment was not available, the next highest resolution taxonomic rank was used to ensure all MAGs were accounted for. Taxa accounting for >5% of the total MAG relative activity in at least one set of sample replicates were plotted on stacked area charts showing average GeTMM relative activity over time separated by treatment. All other taxa were summed into “Other”. **(C)** Conceptual diagram of major prokaryotic community metabolisms in the microcosms as described previously [17]. Previously described primary catechin degraders and secondary consumers of phenolic acids, as well as metabolites previously inferred to move between them, are explicitly labeled. Turquoise arrows specify metabolisms enriched in unamended samples, and orange arrows specify metabolisms enriched in catechin-amended samples.

Given such drastic rewiring of prokaryotic community structure and metabolism, we hypothesized viruses infecting enriched catechin degraders could also impact microbiome community metabolic rewiring. Viruses are already known to be ubiquitous and play an active role across diverse ecosystems, contributing to host mortality and lysis-mediated nutrient release [18] and altering the metabolic objective of host cells during infection (termed virocells) [19]. In soils, there is also mounting evidence viruses influence carbon cycling as inferred from experimental addition of viruses to soils and virus-encoded auxiliary metabolic genes involved in carbon cycling, and linkage of virus dynamics to key carbon cycling host species [20–24].

However, viral response to rapid environmental change, such as abrupt thaw and substrate amendment, and their impact on subsequent community-wide metabolic shifts – intended and unintended – remains virtually unknown. This knowledge gap represents an opportunity to establish baseline data needed to integrate viruses into multi-scale models that predict the fate of carbon in ecosystems worldwide, and critically also capture virus-related unintended consequences to perturbations or engineered interventions [25].

Here, we investigate viral community dynamics and potential functional roles during catechin-mediated methane suppression in permafrost peat microcosms using the same multi-omics dataset from the earlier fen microcosm prokaryote-focused work [17]. Specifically, we leveraged viral identification [26] and host prediction tools [27] to: (i) identify actively transcribed, putatively lytic viruses and their predicted prokaryotic hosts, (ii) quantify viral community responses to catechin amendment across time, and (iii) evaluate evidence for how viruses could impact the microbiome through a viral shunt mechanism that redistributes catechin-derived intermediates to additional hydrogen-consuming polyphenol-degrading taxa, and update our evolving conceptual model for how catechin suppresses methane emissions.

## Materials and Methods

### Sample Collection

See *Peat sampling*, *Incubation set-up,* and *Nucleic acids extraction* in [17] for full description. Briefly, a total of 40 anoxic microcosms were constructed from peat from a fen in Stordalen Mire (Abisko, Sweden) 9-19cm below the surface, where half were left as controls and half were catechin-amended (**Figure 1A**). To generate bulk metagenomes, one microcosm per treatment (catechin, unamended) was sacrificially sampled at days 0, 14, 21, and 35 post-amendment. DNA was extracted and used for Illumina sequencing library construction, which was sequenced up to 500M reads with a median depth of 421M reads. To generate bulk prokaryote metatranscriptome sequencing data, and assess variation, three microcosms per amendment were sacrificed at days 0, 7, 14, 21, and 35 post-amendment. RNA was extracted, rRNA depleted, and DNase treated. The resulting RNA was reverse-transcribed to cDNA, and Illumina sequencing libraries were constructed and sequenced to a depth of up to 100 million reads (median depth: 99 million reads). Day 0 only had metagenomic and metatranscriptomic sequencing for unamended samples because it was sacrificed immediately after the microcosms were sealed and served as the initial timepoint for both treatments. Sequencing sample accession numbers are available in **Supplementary Data 1**.

### Metagenome Assembly and MAG recovery and taxonomic/functional/transcriptomic assessment

See *Metagenome sequencing, assembly, and binning* in [17] for full description. Briefly, metagenome sequencing libraries were assembled with three strategies: (1) Samples individually assembled with MEGAHIT (v1.2.9) [28], (2) Iterative assembly of select individual samples, whereby trimmed metagenome reads from individual samples were mapped against medium and high-quality MAGs generated from the individual assemblies, and reads that did not map were assembled again. Assembled contigs were binned, and the process was repeated until no or few medium or high-quality MAGs were recovered. (3) Co-assembly of different samples with MEGAHIT, where assembled contigs were binned using the combined sample reads to generate coverage profiles. MAGs resulting from the microcosm-derived metagenomes were combined with a set of MAGs that were recovered from a decade of sampling at Stordalen Mire. This full MAG set was dereplicated at 99% ANI, 10% minimum aligned fraction with Drep [28] to produce approximately 2,302 strain-level operational units.

MAGs were then analyzed for taxonomy, functional content, and activity as follows. First, MAG taxonomy was inferred using the Genome Taxonomy Database Toolkit (GTDB-tk v2.3.0 r214) [29]. Second, MAGs were annotated using DRAM (v1.4.4) [30] and CAMPER (v1) [31].

Third, relative expression was approximated with gene length corrected trimmed mean of M-values (GeTMM) [32]. Relative expression of MAG genes were taken from extended data computed previously [17]. MAG relative expression was computed as the average relative expression per gene, calculated by summing the GeTMM values of active genes and dividing by the total number of genes per MAG.

### Virus database construction

Viral contigs were mined from (i) all assembled contigs >5kb from Stordalen peat fen microcosm metagenomic samples [33], (ii) medium and high-quality (est. completeness >50%, <10% contamination) MAGs recovered from the microcosms [17], and (iii) vOTUs previously identified from Stordalen Mire samples [22, 34]. Viral contigs were identified with GeNomad (v1.8.0) [26] using default settings. GeNomad also trimmed host regions from contigs it identified as provirus. Next, contigs considered viral based on GeNomad default post-classification filtering were further quality scored with CheckV (v1.0.3) [35] using default settings, which identified additional provirus sequences, trimmed host regions, estimated viral contig completeness, and estimated contamination from non-viral genes. Viral contigs were then clustered into viral operational taxonomic units (vOTUs) in accordance with the Minimum Information for an Uncultivated Virus Genome (MIUViG) standards [36]. vOTUs were annotated with prodigal-gv (v2.6.3) [37] via the GeNomad annotate module, and inputs for DRAMv functional annotation were prepared with the MVP pipeline (v1.0)[38] module 06. DRAMv (v1.4) [30] was then used to functionally annotate vOTUs.

### Classifying active vOTUs

See *Metatranscriptome Sequencing and Analysis* in [17] for sequencing, read quality filtering, and subsampling procedures. To determine which vOTUs were transcriptionally active, metatranscriptomes were randomly subsampled to 50M pairs of trimmed reads and then aligned to the vOTU reference database using Bowtie2 (v2.5.1) [39] with the following parameters: -D 10 -R 2 -N 1 -L 22 -i S,0,2.50. Output SAM files were converted to sorted BAM files using samtools (v1.18) [40]. Reads mapping <90% average sequence identity to vOTUs were filtered out of the sorted BAM files using bbtools (v38.97) reformat.sh [41] with the parameters: idfilter=0.90 pairedonly=t primaryonly=t. The 90% cutoff was chosen based on viral population read mapping cutoffs established previously [42]. The number of reads mapped to each vOTU gene were counted by first predicting open reading frames with prodigal-gv (v2.6.3), then using htseq-count (v.0.13.5) with the following parameters: -a 0 -t CDS -i ID --stranded=reverse, which filtered out any reads that ambiguously mapped to more than one gene. Read counts <5 were set to zero, as in the previous study [17] and another soil virus metatranscriptomics study [43]. vOTU genes were annotated with DRAMv [30] and GeNomad [26], and database annotations were concatenated into a single string for each gene.

We considered a vOTU conservatively transcriptionally active in a sample if it had (i) ≥5 reads mapping to ≥1 gene that contained any of the substrings “capsid”, “tail”, “package”, “packaging”, “portal”, “lysis”, “terminase”, “spike”, “baseplate”, “internal virion”, “neck”, or “prohead” in its concatenated annotation, a similar strategy to [44] for classifying active-lytic viruses, and (ii) ≥10% of the bases on the viral contig were covered by at least one read using bedtools genomecov [45]. If a vOTU did not meet the requirements of (i) and (ii) in a sample, its coverage was set to zero in that sample. Three additional, more permissive, annotation-independent activity thresholds adapted from previous studies were used to compare results since no benchmarked standard exists: 1) >10% of contig covered by ≥1 read, 2) ≥1 gene with ≥1 read mapped / 10kb, and 3) ≥1 read mapped.

### vOTU characterizations and analyses

#### Activity

The relative expression of vOTUs in metatranscriptomes was calculated using GeTMM normalization method [32] from the EdgeR package (v4.4.2) in R (v4.4). Read counts per vOTU, per sample, that met the criterion above (not set to zero) were GeTMM normalized, then the GeTMM values of each vOTU’s active genes were summed and divided by the total number of genes within the vOTU per sample, yielding an average vOTU GeTMM expression per sample. To determine the rank abundance of vOTUs under different treatments, we identified the maximum GeTMM relative expression attained by each vOTU across all replicate samples within both the catechin-amended and unamended treatments. Species accumulation (collector’s curve), rank abundance, and visualizations were done in Python (v3.12).

#### Alpha and Beta Diversity

vOTU alpha diversity was calculated from active vOTU GeTMM values using first-order Hill numbers (equivalent to the exponential of Shannon H-index) from scikit-bio (v0.6.3) [46] in Python (v3.12). The same method was used to determine MAG alpha diversity. A one-sided Welch’s t-test was used to test the hypothesis that alpha diversity in unamended samples was greater than in catechin-amended samples day 7-35, and that % RNA mapped to vOTUs in unamended samples was less than in catechin-amended samples. Significant changes in value with respect to time were evaluated with a Kruskal-Wallis test. To isolate the treatment effect from temporal dependency, a z-score standardization within each timepoint was applied to center the mean of each timepoint at zero. % RNA mapped values were log-transformed before testing to satisfy the normality assumption. A post hoc power analysis was performed to confirm pooling samples sufficiently increased power over testing differences at individual timepoints (**Supplementary Data 3**).

vOTU beta diversity was determined using Hellinger-transformed Bray-Curtis dissimilarity of vOTU GeTMM values using vegan (v2.6-10) [47] in R. Beta diversity was visualized using principal component analysis. The same approach was used to calculate and visualize MAG beta diversity. Three sample clusters were designated based on the visual clustering of samples on the PCA and response category: 1) *shared early response*: catechin day 7 and unamended day 7 samples, 2) *catechin response*: catechin day 14, 21, and 35 samples, and 3) *unamended response*: unamended day 14, 21, and 35 samples. A global PERMANOVA was used to test if the centroids and/or dispersion between the three sample clusters were significant, as well as pairwise PERMANOVA between cluster pairs. To test if Bray-Curtis distances between treatments were significantly different across days, we performed pairwise Wilcoxon rank sum tests and visualized significant differences with compact letter display using multCompView v1.9-1 in R.

### Virus-host prediction

Virus-host predictions were made using iPHoP (v1.3.2) [27]. A custom iPHoP host database was constructed by adding the 2,302 MAGs identified from microcosm samples and previous Stordalen analyses to the iPHoP standard database (iPHoP_db_Aug23_rw release). iPHoP was run on vOTU sequences using the custom database and default settings, with vOTU-MAG predictions available in **Supplementary Data 4**. vOTU-MAG predictions were visualized with Cytoscape [48], and metadata from the node table available in **Supplementary Data 5**. Predictions between active vOTUs and active MAGs were considered first, and additional host predictions were made based on active vOTUs predicted to infect MAGs from the iPHoP standard database or inactive bioreactor MAGs. Another set of host predictions were made based on gene-sharing networks, whereby vOTUs in the same taxonomic cluster without a host assignment were putatively assigned if all other vOTUs in the cluster were assigned to the same host genus. Taxonomic clusters were generated with vConTACT3 v3.0.5 using database v228 and default settings, with results available at **Supplementary Data 6**.

### Auxiliary metabolic gene (AMG) identification

AMGs were identified from active vOTUs using DRAMv annotations. A gene was considered an auxiliary metabolic gene if it contained an “M” (metabolic) flag, but no “B” (consecutive metabolic genes) or “F” (near-end-of-contig) flags, and was both upstream and downstream of a gene with a “V” (viral) flag [30, 49, 50]. All detected AMG annotations are available in **Supplementary Data 7**.

### Indicator Species Analysis

Indicator values were computed per vOTU using GeTMM values. The following five clusters were used as the sample groups to compare vOTU relative abundance between: unamended day 0, unamended and catechin day 7, unamended day 14-35, catechin day 14 and 35, and catechin day 21. Indicator values were computed using the multipatt function from the indicspecies package in R with parameter duleg=False to consider combinations of site groups [51]. vOTUs with a p-value < 0.01 were classified as indicators (**Supplementary Data 8**).

### Metagenomic relative abundance comparisons between viruses and their predicted hosts

Metagenomic samples were mapped back to vOTUs and MAGs with Bowtie2 (v2.5.1) [39] with the following parameters: -D 10 -R 2 -N 1 -L 22 -i S,0,2.50. Output SAM files were converted to sorted BAM files using samtools [40]. Relative abundance was determined using CoverM [52] with the following flags for vOTUs: mode=trimmed_mean, min-read-percent-identity=90, min-alignment-fraction=75, min-covered-fraction=10, and for MAGs: mode=mean, min-read-percent-identity=97, min-alignment-fraction=75, min-covered-fraction=75.

### Comparative genomics and transcriptome coverage analysis

Active vOTUs predicted to infect MAGs in the genus JAGFXR01 were clustered in VirClust with default settings [53]. Briefly, open reading frames were predicted using MetaGeneAnnotator [54], and compared using all-vs-all BLASTP. Then, genome distances were calculated using markov clustering. Read pileup was calculated for contig_591846 in catechin-amended samples using samtools mpileup based on the filtered, sorted BAM files generated from metatranscriptomic samples. Contig_591846 genes were visualized with gggenomes [55], and read pileups were visualized with ggridges [56] in R. Contig_591846 was aligned to JAGFXR01 MAG STM_0716_E_M_E034_A_bin.10 with NCBI BLAST blastn on the NCBI web portal using default parameters and visualized in R.

## Results + Discussion

### Experimental design, dataset overview, and virus recovery

Our prior work leveraged climate-sensitive peat microcosms from a Stordalen Mire fen to evaluate catechin-suppressed methane production from the prokaryotic community perspective [17]. Catechin amendment decreased methane emissions by >80% from the fen microcosms.

To uncover the mechanism/s underlying this inhibition, bulk metagenomes, metatranscriptomes, and metabolomes were obtained from the microcosms along a 35 day time course (**Figure 1A**, see Methods for details) and analyzed for metabolic signal. These past MAG-resolved metatranscriptome data analyses revealed prokaryotic community expression was dominated by *Clostridium* and *Pseudomonas_E* in nearly all samples, with *Paludibacter* enriched in unamended samples and undescribed *Bacillota* genus JAGFXR01 enriched in catechin-amended samples (**Figure 1B**). These dominant taxa played key roles in unamended and catechin-amended carbon metabolism (**Figure 1C**), namely polysaccharide metabolism and catechin and phenolic acid metabolism, respectively. Together these findings revealed a microbial community-wide metabolic transition from a syntrophic network that feeds methanogenesis to a catechin fermentation network. Here we evaluate if, and to what extent, viruses play a role in this community-wide metabolic transition by identifying actively lytic viruses, predicting their hosts, and investigating how viral communities responded to catechin amendment in the context of diverse biomolecular measurements.

We first established relevant reference genome resources by identifying virus contigs from microcosm metagenome assemblies (this study) using GeNomad [26], a state-of-the-art virus identification tool with a reported accuracy of >94% for contigs >5kb, which yielded 21,882 virus contigs >5kb. These contigs were clustered with 6,645 DNA viral operational taxonomic units (vOTUs) derived from previous virus analyses in Stordalen Mire [22, 34] for a total of 9,562 vOTUs, where an additional 3,994 unique vOTUs were identified from this study (**Figure S1A**).

From these, we next assessed which vOTUs were active in at least one microcosm metatranscriptome based on activity thresholds. Because no established standard exists for defining viral activity from metatranscriptome data [57], we employed four different activity thresholds, which resulted in as little as 900 to as much as 3,772 of the 9,562 vOTUs being considered as active (**Figure S1A, S2A,** see methods), and since it is unknown where in this spectrum of conservative to permissive is most correct, we present the most conservative findings in the main figures of this study and interpret results within the context of these thresholds when relevant. Operationally, the most conservative activity threshold required (i) ≥5 reads mapped to an annotated gene involved in viral particle formation or lysis, which is the median number of reads mapped to genes with ≥1 read map and the same strategy to determine gene activity as in the previous study [17] and (ii) >10% of bases on the viral contig were covered by at least one read. These thresholds ensured that viruses classified as active were transcribing genes associated with viral particle formation or lysis, a hallmark of lytic activity [58], and decreased false positive activity classification from non-specific read mapping by requiring a minimum coverage breadth [59]. Additionally, we explored three more permissive, annotation-independent thresholds (≥10% of contig mapped by ≥1 read, ≥ 1 gene mapped by ≥1 read / 10kb, and ≥1 read mapped) (**Figure S2**) to evaluate robustness and as a control for the possibility that the most conservative threshold may remove poorly annotated and/or lowly active viruses.

Towards interpretation, we used the most conservative threshold for the main results as this represented the most confident set of lytic active vOTUs since they contained verifiably transcribed late-stage infection viral genes. However, caution is still warranted. This is because, even though GeNomad is highly accurate (>94% MCC for contigs >3kb [26]) and our conservative activity threshold would filter out any potential mobile genetic elements that did not contain viral particle formation or lysis gene annotations, our method may retain other virus-related biological entities (i.e., gene transfer agents [60] and contractile injection systems [61]). The prevalence of gene transfer agents in environmental samples is unknown as only a few have ever been cultured, and systematically identifying them bioinformatically and experimentally continues to be a grand challenge [60]. As for contractile injection systems, these primarily encode cellular Type VI secretion systems and were likely included in the chromosomal dataset when GeNomad was trained, but others like tailocins are derived from defective prophages and may be indistinguishable from transcribed virus tail fragments [62].

Even considering only the conservatively active vOTUs, sampling curves suggested that active vOTUs were relatively well sampled in the metatranscriptomes (**Figure S1C, S2B**) and rank abundance analyses suggested that even lowly active vOTUs were captured (**Figure S1D, Figure S2A**).

### Viral community activity responses differ across treatments

Given well sampled active vOTUs, we next sought to evaluate the catechin response across the experimental duration (days 7-35) by pooling samples into unamended (n=11) and catechin-amended (n=12) groups. A post-hoc power analysis confirmed that this sample size provided 97.7% power to detect significant differences at alpha = 0.05, which was significantly more powerful than testing significance at individual timepoints (**Supplementary Data 3**). Because alpha diversity shifted significantly over time (Kruskal-Wallis, *P* = 0.0445), we applied z-score standardization within each timepoint to isolate the treatment effect from temporal variance. This transformation successfully neutralized time-based bias, as confirmed by a subsequent Kruskal-Wallis test on the z-scores (*P* = 0.99). Following normalization, the data met the assumptions for parametric testing (Shapiro-Wilk, *P* > 0.05). A one-tailed Welch’s t-test revealed that catechin amendment triggered a significant reduction in vOTU alpha diversity compared to unamended controls at all activity thresholds except >10% of contig mapped (**Figure 2A, S2C,** upper panels). We observed an inverse trend in the proportion of RNA mapped to active vOTUs (**Figure 2A, S2C**, lower panels). After log-transformation and z-score standardization to satisfy normality and remove the time bias, respectively, Welch’s t-test confirmed that catechin amendment significantly increased the relative proportion of mapped viral RNA across all activity thresholds (*P* < 0.01). Together, these results support that catechin amendment triggered a specific relative increase in lytic virus activity within a subset of the viral community, and significantly decreased alpha diversity in three of the four activity thresholds tested.

**Figure 2.**
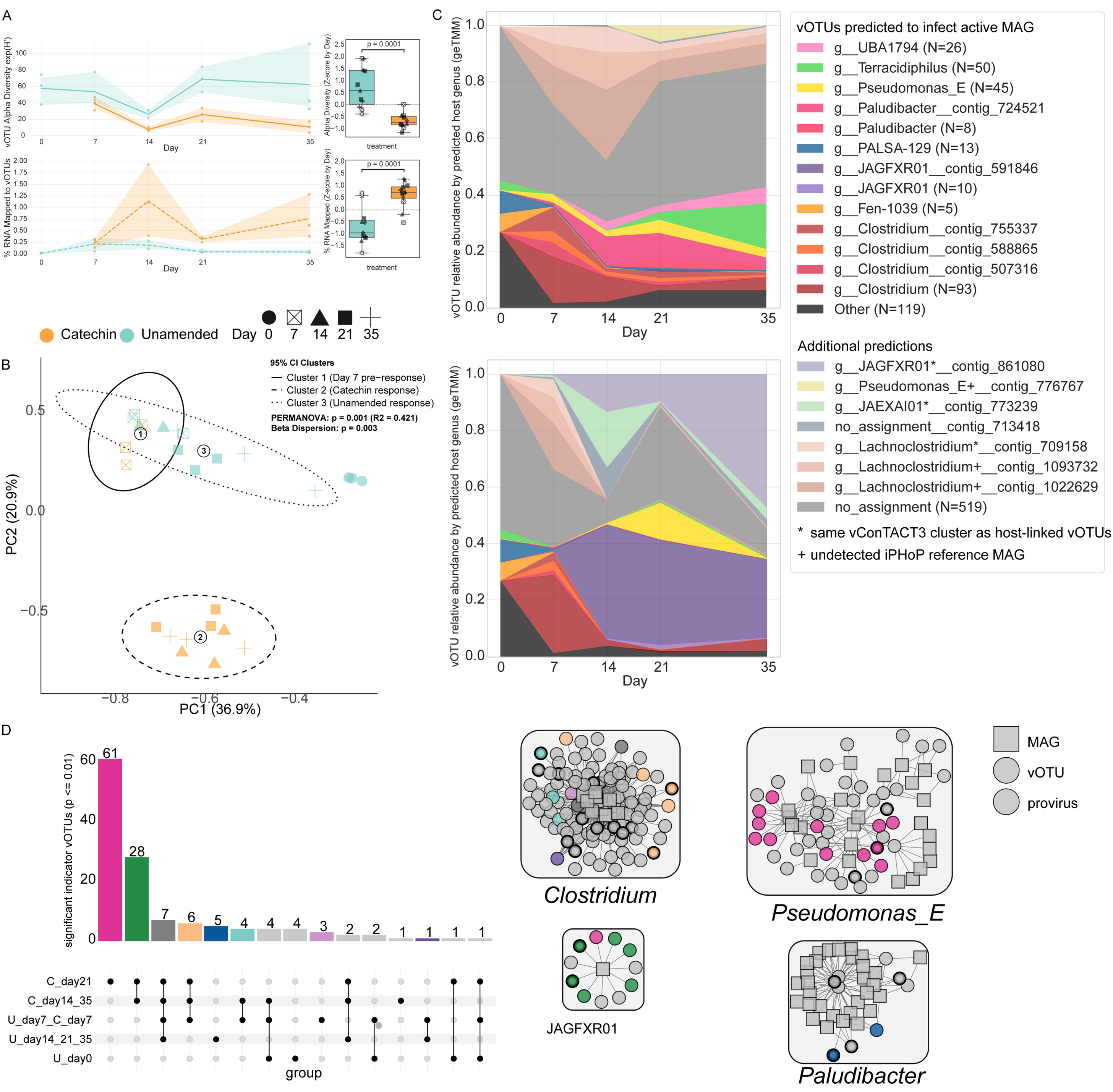
vOTU communities differed between treatments and over time. **(A)** vOTU alpha diversity (Hill Number of order = 1) and % of total metatranscriptomic reads mapped to active vOTUs. Shaded areas represent 95% confidence intervals. Boxplots show distribution of alpha diversity per treatment, with unique shapes representing different timepoints. Values were z-score standardized within each timepoint to center the means and remove temporal autocorrelation. *P*-values are based on one-sided Welch’s t-tests on z-scored alpha diversity values. **(B)** Principal component analysis of Bray–Curtis dissimilarity among metatranscriptomic samples using active vOTU GeTMM relative abundance values. Samples labeled by day; colors correspond to treatment. Black circled numbers represent centroids of sample clusters. Dashed lines represent 95% confidence intervals. PERMANOVA and Beta Dispersion results are based on a global test including all three clusters. **(C)** Stacked area chart of active vOTU GeTMM relative activity in unamended samples (top) and catechin-amended samples (bottom) aggregated by predicted host genus (sum). Aggregated genera with <5% contribution in at least one set of sample replicates were grouped into “Other”. vOTUs accounting for >4% of average GeTMM relative activity in at least one set of sample replicates were plotted individually in a similar shade to their predicted host genus. Host predictions are labeled by their supporting analyses. **(D)** (left) Upset plot of indicator species analysis of active vOTUs across sample clusters (p-value ≤ 0.01). Upset plot y-axis is labeled by which samples were assigned to each group. All replicates of the same timepoint were assigned to the same group. (right) Infection network of key carbon cyclers. Squares represent MAGs, circles represent vOTUs, and bold circles represent proviruses. Edges represent iPHoP phage-host predictions with confidence score >90. The circle color denotes the combination of sample clusters the vOTU was an indicator for and corresponds to the bar colors in the upset plot.

Next, we investigated beta diversity to determine how similar viral community expression was across treatments and timepoints (**Figure 2B, S2D**). From the initial timepoint to day 7, both catechin and unamended vOTU community expression shifted considerably from the day 0 timepoint (**Figure S3,** average Bray-Curtis distance = 0.859, ± 0.054 SD); however, community structure remained relatively similar between treatments at this stage (average Bray-Curtis distance = 0.492, ± 0.092 SD), and were not significantly different (PERMANOVA: R^2^ = 0.45, *P* = 0.1). This shift was observed across all activity thresholds and suggests early changes in vOTU community expression presumably reflected bottle effects as the communities acclimated to the bioreactor environment rather than a treatment effect. However, vOTU community expression in catechin-amended samples days 14-35 diverged significantly from day 7 across all activity thresholds (PERMANOVA: *P*_adj_ < 0.01). Even though beta dispersion was significantly different between sample clusters (Wilcoxon: *P* < 0.05), visual separation of clusters and lack of overlap between confidence intervals on the PCA support that catechin day 14-35 samples represented a distinct shift in vOTU community expression, and more variable than day 7 samples. These catechin-amended samples also differed significantly from unamended samples day 14-35 (*P*_adj_ < 0.01) and beta dispersion was insignificant (Wilcoxon: *P* > 0.05), which reinforced that catechin treatment enriched for a distinct community of viruses. While unamended samples day 14-35 also shifted from day 7 (*P*_adj_ < 0.01), they maintained closer proximity to the day 7 state, as evidenced by overlapping confidence intervals. Collectively, these results demonstrate that catechin amendment enriched a unique subset of vOTUs, with the treatment effect becoming prominent by day 14. Additionally, community-level ecological patterns still emerged despite high turnover in active vOTUs between samples, even when using the most conservative activity threshold, where the average vOTU was active in only two samples (**Figure S1E**), similar to previous observations in soil virus environmental studies [34, 63, 64].

We wondered how these viral community diversity patterns compared to those derived from the MAGs captured in these data – observations not previously reported in our earlier work and therefore new to this study. These analyses revealed that, similar to vOTU alpha diversity, expression-based MAG alpha diversity was significantly lower in catechin-amended samples than in unamended samples (**Figure S4A**, *P* < 0.01). Beta diversity of prokaryotic community expression also followed similar patterns to vOTU community shifts: a clear departure from the initial timepoint by day 7 followed by distinct catechin and unamended community responses, with the unamended response more similar to the initial timepoint (**Figure S4B**, PERMANOVA: *P* ≤ 0.001). To determine if MAG community activity changed more gradually compared to vOTU community activity at the most conservative activity threshold, Bray-Curtis dissimilarity between samples of the same day but different treatments were compared (**Figure S4C**). MAG community activity overall was more similar between treatments, and Wilcoxon rank sum tests supported that MAG community activity between treatments was significantly more diverged at day 35 than at day 7, but that the divergence was gradual through day 14 and 21. In contrast, vOTU community activity by day 14 was significantly more different between treatments than by day 7, but we did not observe vOTU community activity become significantly more different by day 21 or 35. These different patterns suggest that vOTU community expression changes were not strictly coupled to the MAG community expression, and vOTU community expression changed more dramatically than the MAG community over time and across treatments. We posit that this delayed viral activity response stemmed from the relatively small day 7 prokaryote differences, which may have been insufficient to induce a viral community-level shift in activity.

Previous perturbation studies in rewetted soils demonstrated a subset of responding vOTUs dominated viral community activity, in sync with enriched taxa that were predicted to be hosts [63, 65]. Given these findings, and our observations above, we hypothesized a subset of newly dominant viruses may have driven the catechin-amended treatment shift in viral community activity from day 7 to 14 samples, presumably by infecting previously identified catechin-enriched microbes (**Figure 1B**).

### Viral community expression was dominated by vOTUs predicted to infect key carbon cyclers

We predicted hosts for active vOTUs using a database augmented with sample-specific MAGs and iPHoP, a machine learning based host prediction tool that aggregates many *in silico* approaches into a single probabilistically modeled prediction [27]. This generated an infection network (**Supplemental Data 5 & 6**) derived from 41.5% (374/900) of active vOTUs with host predictions, 66% (249/374) of which were based on sequence homology (i.e., BLAST, see Supplemental Data 4), spanning 56 transcriptionally active genera and 13 phyla. This fraction of vOTUs that could be assigned a host more than doubles that in other non-human associated microbiome studies (15-22%) [27, 34] and is on-par with recent soil virus studies (37.8-44.5%) [44, 66]. With these host predictions, we first explored whether any viruses were predicted to infect methanogens, which might influence methane metabolism through (i) increased viral activity coupled with decreased methanogen MAG activity, or (ii) auxiliary metabolic gene expression. While few methanogen viruses have been experimentally identified [67], recent studies have attempted to identify them with CRISPR spacers and host-derived auxiliary metabolic genes [68, 69], and hinted at their potential to impact methanogen metabolism [69]. In our study, conservatively active vOTUs were predicted to infect *Methanothrix*, *Methanosarcina,* and *Methanobacterium_A* (**Figure S5A**), and were generally more active in unamended vs catechin-amended samples (**Figure S5B**). Despite detecting actively lytic vOTUs predicted to infect methanogens, no clear treatment-specific infection dynamics were observed. This absence of discernible patterns may reflect the relatively low abundance or slow growth, manifest in sparse metatranscriptome sequencing coverage, of methanogens and their viruses within the community. Additionally, we did not detect any auxiliary metabolic genes within these vOTUs (**Supplementary Data 6**). Overall, any impacts methanogen viruses might have had were not detectable in our analyses from these bulk metatranscriptomic data.

Beyond methanogens, we did observe that, on average, 67.2% (± 16.8% SD) of viral community expression could be linked to an active MAG at the most conservative activity threshold, and an even greater proportion at more permissive activity thresholds. This signal was dominated by vOTUs predicted to infect other key carbon cyclers, with distinct responses observed in catechin-amended samples (**Figure 2C, S2D**). In unamended samples, the bulk of viral community expression was from vOTUs predicted to infect *Clostridium* and genera previously identified as catechin-sensitive [17] (i.e., prokaryotes whose abundance decreased following catechin amendment) such as *Paludibacter*. In contrast, viral community expression in catechin-amended samples was dominated by viruses predicted to infect genera previously identified as key members of catechin degradation processes, namely undescribed *Bacillota* genus JAGFXR01, *Clostridium*, and *Pseudomonas_E* [17] (**Figure 1B**). Both treatments showed Clostridium vOTU dominance at day 7, but diverged by day 14. JAGFXR01-infecting vOTUs surged in catechin-amended samples by day 14 and remained elevated, while *Pseudomonas_E* viruses showed a transient bloom at day 21. These patterns mirror those observed in the initial diversity analyses (**Figure 2A, 2B**) with nearly all of the four-fold increase in the metatranscriptomic read mappings of detected viruses from day 7 to 14 (**Figure 2A**) attributable to predicted JAGFXR01-infecting vOTUs (**Figure S6**).

Several conservatively active vOTUs were highly expressed in specific treatments, suggesting potential ecological importance, but lacked active host MAG predictions (**Figure 2C**). These vOTUs either were predicted to infect inactive MAGs recovered from the bioreactor, iPHoP reference database MAGs, or lacked relevant genetic information to make a confident prediction. These vOTUs might also be genetically related to other recovered vOTUs that did have host predictions, and while genetic similarity does not mean they infect the same host [70, 71], it has been observed for some lineages that viruses and their hosts co-evolve, and viruses that infect the same host may share genes more frequently with each other than viruses that infect different hosts [72, 73]. Therefore, host predictions for vOTUs genetically related to the highly active vOTUs without host predictions may give an idea about their potential host target. Based on this, we explored the potential hosts of these highly active vOTUs through (i) predictions to inactive or reference MAGs in the iPHoP host database, and (ii) taxonomic similarity through gene-sharing networks (see Methods) to other vOTUs with a host prediction. Towards the first approach, the two most abundant unassigned vOTUs in unamended samples were predicted to infect *Lachnoclostridium phytofermentans* reference MAGs, which was reclassified as *Clostridium* [74], and another previously unassigned vOTU was predicted to infect *Pseudomonas_E* reference MAGs (**Supplementary Data 4, Figure S7**). Towards the second approach, all but one unassigned vOTU clustered with at least one other vOTU recovered in this study with a host prediction. Within each vOTU infection network, all vOTUs with a host prediction were predicted to infect the same host genus (**Figure S7**), of which *Pseudomonas_E*, JAGFXR01, and JAEXAI01 were previously identified as polyphenol degrading, catechin-enriched taxa [17]. All additional assignments were to key carbon cyclers previously identified. We included these assignments as “Additional Predictions” annotated with the origin of the prediction. Interpreting these additional assignments with the iPHoP assignments point towards *Clostridium*-infecting viruses as the dominant (>50%) active virus group at unamended day 7 and 14, and catechin-amended day 7 samples, and JAGFXR01-infecting viruses dominating in catechin day 14-35 samples.

Together, these relationships suggest these highly active vOTUs infected ecologically relevant hosts and that viral predation of key carbon cyclers may be even higher than predicted above. While viral taxonomy is not an explicit feature used in virus-host prediction algorithms, future methods could benchmark the usefulness of viral taxonomy in virus-host predictions at different taxonomic ranks.

### Differential virus responses within key carbon cycler infection networks suggests vOTU-specific niches

A subset of MAGs from each genus of key carbon cyclers responded to catechin amendment [17], implying approximately strain-level differential responses that might correspond to distinct niches, as observed in other systems [75, 76]. Thus, we posited vOTUs predicted to infect these key carbon cyclers would also differentially respond, indicative of distinct vOTU-level niches [77, 78]. To initially filter for the subset of responding vOTUs, we employed indicator species analysis [51] as a statistical framework to determine which vOTUs were strongly associated with specific sample clusters. We chose five sample clusters based on PCA of Bray-Curtis distances (**Figure 2B**) and relative abundance analysis (**Figure 2C**): the initial timepoint day 0, the vOTU community acclimating to bioreactor conditions before treatment response (unamended and catechin day 7), the unamended response (unamended day 14-35), the catechin response dominated by vOTUs predicted to infect JAGFXR01 (catechin day 14 and 35), and the catechin response where a transient bloom in vOTUs predicted to infect *Pseudomonas_E* and unassigned vOTUs is observed (catechin day 21, **Figure 2C**). Indicator species analysis revealed 130 indicator vOTUs for at least one combination of sample clusters (*P* value ≤ 0.01, **Figure 2D**). Within the key carbon cycler infection networks, we observed indicator vOTUs for five (for *Clostridium*), one (for *Pseudomonas_E*), two (for JAGFXR01), and one (for *Paludibacter*) different sample cluster combinations, demonstrating differential responses between treatments and/or over time within infection networks (**Figure 2D**). Specific niches were most apparent in vOTUs predicted to infect *Clostridium*, and potentially reflect the diversity of strain-level responses within *Clostridium* hosts. These results support that a subset of vOTUs predicted to infect key carbon cyclers were significantly enriched in particular samples and suggest existence of different vOTU-specific niches within genus-level infection networks.

Given such varied sub-genus-level vOTU and MAG responses, we explored whether ecological patterning could refine virus-host predictions below the predicted genus level. Virus-host linkages at genus and species ranks are known to be challenging as genome-based prediction methods (e.g., iPHoP [27]) differed greatly compared to experimentally benchmarked approaches such as proximity ligation [79]. Correlation-based inference is also challenging in complex communities, particularly on time-series data [80]. However, we reasoned that coupling genome-based host predictions and correlated co-occurrence dynamics might provide independent evidence to 1) evaluate if vOTUs were significantly more correlated with predicted MAGs than non-predicted MAGs within a genus and 2) distinguish active infections among multiple potential hosts at the species level — at least for the subset of taxa where time-resolved patterning was distinct across time points and/or treatments (**Figure 3A**). Using this rationale, we then tested whether treatment-enriched indicator vOTUs showed correlated dynamics with co-enriched predicted hosts, focusing on the four key carbon cycler infection networks identified above.

**Figure 3.**
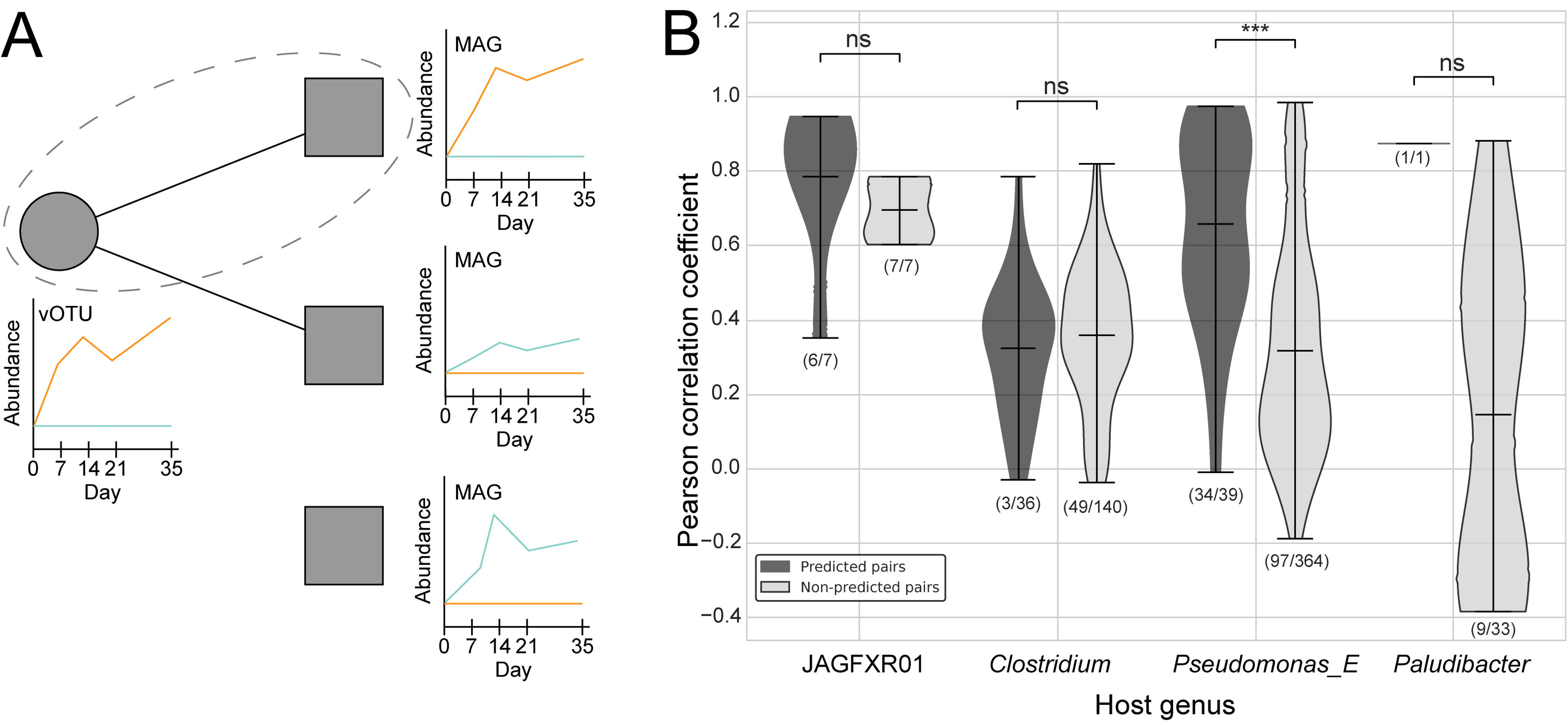
Correlation inference of indicator vOTUs and predicted hosts within key carbon cycler infection networks. **(A)** Conceptual overview of correlation-based inference from metatranscriptomic data. The top virus-host pair demonstrates a virus-host prediction (black line) where vOTU (circle) and MAG (square) relative abundance are correlated across samples. The middle virus-host pair represents an in silico prediction, but where the vOTU and MAG are not correlated. The last MAG is in the same genus as the other two, but the vOTU is not predicted to infect it. **(B)** Violin plot of pearson correlation coefficients between indicator vOTUs and their predicted hosts and non-predicted hosts within each genus-level infection network (from Figure 2D). ns = not significant, *** = *P* value < 0.005. Dark grey plots represent values between iPHoP-predicted vOTU-MAG pairs, and light grey plots represent values between vOTUs and MAGs they were not predicted to infect, but were in the same genus as their predicted host MAG. Values at the bottom of each violin plot represent how many vOTU-MAG pairs were significantly correlated (BH-adjusted *P* value < 0.05) between either predicted MAGs or non-predicted MAGs in the infection network out of the total pairs making up the distribution.

In three of the four key carbon cycler infection networks (JAGFXR01, *Pseudomonas_E*, and *Paludibacter*), nearly all indicator vOTUs were significantly correlated with their predicted hosts based on relative expression across all samples (**Figure 3B**, *P_adj_* < 0.05), qualitatively supporting the MAG-level host predictions within each genus. However, of the four genera tested, only *Pseudomonas_E*-infecting indicator vOTUs were significantly more correlated with predicted MAGs versus non-predicted MAGs in the same host genus, likely because many MAGs within each genus responded similarly across samples (**Figure S8, dashed lines**).

*Clostridium*-infecting vOTUs were only weakly correlated to their predicted host MAGs, and correlation distributions were nearly identical between predicted and non-predicted *Clostridium* MAGs and predicted vOTUs (**Figure 3B**). These results largely refute our first hypothesis that vOTUs would be more significantly correlated with predicted MAGs than non-predicted MAGs within a genus. Yet, within each set of vOTU-MAG predictions for the four key carbon cyclers minus *Clostridium*, indicator vOTUs were significantly correlated with at least one predicted host MAG (**Figure S9**). These observations could indicate that among multiple MAGs within a genus predicted as potential hosts for a treatment-enriched vOTU, co-enriched MAGs may represent the primary targets sustaining viral population activity, though infection of other predicted hosts cannot be excluded. While previous paired virus-prokaryote eco-genomics studies in soil ecosystems, including at this site, have documented correlations between vOTU abundance and their hosts across samples [81], our approach extends this framework by seeking to resolve sub-genus vOTU-MAG relationships. If future experimental measurements can be developed to confirm these relationships, such analytics would allow for community-wide increasing precision for inferring active infection networks within complex systems.

### One dominant vOTU may facilitate rewiring of community carbon metabolism

One of the indicator vOTUs, contig_591846, was predicted to infect the most abundant JAGFXR01 MAG. This prediction was based on BLAST sequence alignment (confidence score: 93.3/100), with 13 hits of an average length of 261 bases ± 168.4. Twelve of thirteen hits returned e-values < 1e-05 (trimmed mean: 6.09e-22 ± 2.02e-21), suggesting recent horizontal gene transfer between the phage and host genomes. Although iPHoP can predict broader host ranges, contig_591846 was exclusively predicted to infect the most abundant JAGFXR01 MAG across iPHoP classifier sub-modules and maintained low prediction scores for the less abundant JAGFXR01 MAG (**Supplementary Data 8**), supporting the high specificity of the BLAST method and confidence in this putative phage-host linkage, and an instance of strain-level host prediction.

Genomically, contig_591846 was (i) high-quality (40 kb, >90% estimated completeness), but not complete (no direct terminal repeats or circularization), (ii) appeared to be a double-stranded DNA, encapsidated, tailed, temperate phage (**Figure 4B**, lower panel), and (iii) shared less than 10% of its proteins with other JAGFXR01-infecting vOTUs (**Figure S10**).

**Figure 4.**
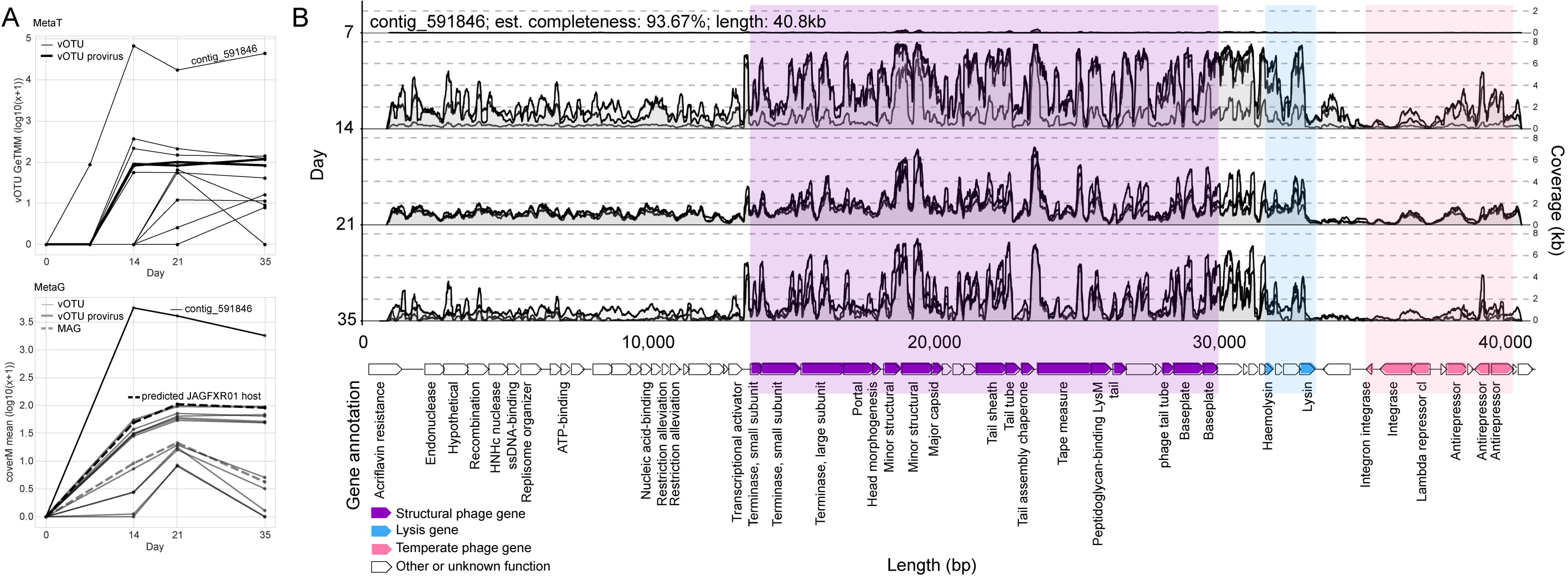
Relative abundance of JAGFXR01 MAGs and predicted viruses. **(A)** GeTMM values of active vOTUs predicted to infect JAGFXR01 averaged across replicate catechin-amended samples (top). Lines are colored by indicator value as in Figure 2D. CoverM trimmed mean coverage of active JAGFXR01-infecting vOTUs and JAGFXR01 MAGs (bottom). **(B)** Contig_591846 read pileup from catechin-amended metatranscriptomic sample reads mapping >90% ANI to the coding strand. Replicate samples are plotted on top of each other. Predicted ORFs are plotted below, with abbreviated functional annotations based on DRAMv and GeNomad. Select genes are colored by function with functional gene regions shaded in similar colors.

Taxonomically, gene-sharing analyses against vConTACT3 [82] showed that contig_591846 clustered with eight viral sequences at the family-level (an unnamed family within phylum *Caudoviracetes*), six of which were RefSeq viruses belonging to genera *Abouovirus* and *Jimmervirus*, and two vOTUs identified in this study (**Supplementary Data 9**). In the microcosm data, this same vOTU was a dominant responder as it accounted for 27.8-40.2% of the overall viral community expression in catechin day 14-35 samples, and 36.8-76.6% of total detected JAGFXR01-infecting vOTU activity (**Figure 2C**, dark purple). Further, its expression was on average 82-308x higher than that of the next most highly expressed vOTU longer than 10kb predicted to infect JAGFXR01 (**Figure 4A**, upper panel). Together these findings support that the dominantly responding contig_591846 was near-complete, genetically divergent from previously identified viruses, and likely an induced prophage lytically responding to the catechin-enriched JAGFXR01 niche.

However, one of the taxonomically related vOTUs identified here, represented by contig_861080, was a short genome fragment (5.4kb, 11.9% CheckV estimated completeness) that was 96.61% similar based on nucleotide identity to contig_591846 and classified in the same subfamily but different genus. Contig_861080 accounted for 20.1–62.6% of the overall vOTU activity in catechin day 14-35 samples, but its limited size precluded confident conclusions regarding its host. Given the high degree of similarity to contig_591846, it’s possible this fragment represents the same virus population, and that JAGFXR01 vOTUs made up nearly 80% of virus community expression by day 35 in catechin-amended samples. Yet, it’s also possible contig_861080 represents an equally dominant, distinct virus population that infects JAGFXR01, or a virus that infects a different host taxa.

To determine if contig_591846 was undergoing induced lysis, we assessed its metatranscriptome expression of functionally relevant lysis and lysogeny genes from days 14, 21, and 35 catechin-amended samples, and its genomic copy number compared to that of its predicted JAGFXR01 host MAG. Metatranscriptome analysis revealed that contig_591846 in catechin-amended samples (i) consistently expressed structural and lysis gene regions, (ii) expressed the integrase gene region, although it was two- to three-fold lower than expression in the structural gene region, and (iii) antirepressor gene expression was two- to three-fold higher than that of the cI repressor across catechin-amended samples (**Figure 4B**). These observations are consistent with signature gene expression expectations for lytically-active phages [83, 84], and that, perhaps, a subset of the viral population was also remaining lysogenic as a non-induced prophage. Metagenome analysis revealed genomic representation (where metagenomes were available) that ranged from 20- to 156-fold more abundant than its predicted JAGFXR01 host MAG (**Figure 4A**, lower panel), indicative of the sustained presence of virus genomic copies far exceeding prophage-only abundances and indicative of at least a subset of the virus population replicating its genome during lytic infection.

Metagenome-based virus-host microbe ratios vary widely [85], and are not typically reported at the vOTU-MAG level due to prediction tool imprecision (discussed above). Thus, we treated this unique treatment-responder situation as an opportunity to push biological inferences, with a focus on attempting to genomically assess the average per-cell viral burst size for this lineage. Specifically, JAGFXR01 vOTU-MAG relative abundance ratios >100 from days 14-35 suggest a bust size of at least this (depending upon the proportion of the population remaining as prophage), which we posit to indicate that JAGFXR01 is undergoing intense, sustained viral replication.

Given this evidence for intensive lytic infection pressure, we wondered what downstream effects JAGFXR01 lysis might have on the broader microbial community. Previous soil virus studies experimentally demonstrated that viruses alleviate some microbial nutrient limitations [86], perhaps through frequently lysing abundant hosts to increase diversity via auxiliary community members scavenging nutrient-rich lysates. In our system, we reasoned that enrichment of catechin-degrading JAGFXR01 cells would open a new niche for opportunistic viruses, generating a “kill-the-winner” [87] scenario for responding viruses, where viral predation rapidly increases for fast-growing hosts as a top-down control, and this in turn could stimulate community diversity via lysis-released partially metabolized catechin intermediates diversifying resources available to additional polyphenol-degrading microbes. Indeed, we previously showed aromatic catechin degradation intermediates accumulated in catechin-amended sample pore waters, and that hundreds of additional MAGs expressed genes for polyphenol metabolism but not catechin degradation [17]. However, viral lysis was not considered and it remained unclear if these non-catechin-degrader phenol metabolisms were enriched in catechin-amended samples.

To investigate whether additional polyphenol-degrading MAGs were utilizing these downstream intermediates and if these metabolisms were enriched in catechin-amended samples, we assessed the relative expression of genes encoding key catechin-degradation intermediate metabolism (phloroglucinol transformation genes *PGR*, *pgthAB*) in 104 medium-and high-quality MAGs that lacked catechin-specific gene annotations. Compared to unamended samples, this revealed significantly higher cumulative relative expression of *pgthAB* genes from these MAGs in all 3 virus-active (catechin-amended days 14-35) time points (**Figure S11A**, Welch’s t-test: *P* = 0.00129). However, *PGR* expression was not significantly different (**Figure S11B**, Welch’s t-test: *P* = 0.209). Individually, of the 104 MAGs expressing genes for aromatic intermediate metabolism but not catechin degradation, a total of 34 MAGs drove the higher expression with 25 MAGs (representing 21 genera), 7 MAGs (5 genera), and 2 MAGs (2 genera) showing elevated expression of at least one phloroglucinol transformation gene in catechin-amended samples at 1, 2, or 3 timepoint(s), respectively during the virus-active period (**Figure 5A**). Further, phloroglucinol degradation requires hydrogen, and hydrogen-uptake hydrogenase activity among these MAGs was significantly elevated in catechin-amended samples day 14-35 (**Figure 5B** *P* < 0.0001). In fact, nearly all (94.5%) of hydrogen-uptake hydrogenase expression was attributable to two undescribed *Actinomycetes*, JAEXAI01 and JAATFL01. While other *Actinomycetota* contained catechin-specific gene annotations, these two did not contain such genes in the recovered MAGs and were not explicitly mentioned in the previous study. These additional polyphenol degraders accounted for 78.1% and 16.4% of hydrogen-uptake hydrogenase expression, respectively, even when representing only 2–4% and 0.54–2.6% of total relative genome-wide MAG activity in catechin-amended samples (days 14-35). Although JAEXAI01 (CheckM2 97.72% complete, 0.64% contamination) and JAATFL01 (CheckM2 78.71% complete, 0.14% contamination) lacked specific annotations for flavonoid degradation, both possessed genes for lignan, phenolic acid, pyrogallol, and phloroglucinol degradation, which were nearly all significantly more active in catechin-amended samples than in unamended controls (**Figure S12**, Welch’s t-test: *P_adj_* < 0.05) and lead us to hypothesize that the primary mechanism of catechin amendment—suppression of methanogenesis via competition for hydrogen--might be predominantly driven by these two additional, previously undescribed MAGs.

**Figure 5.**
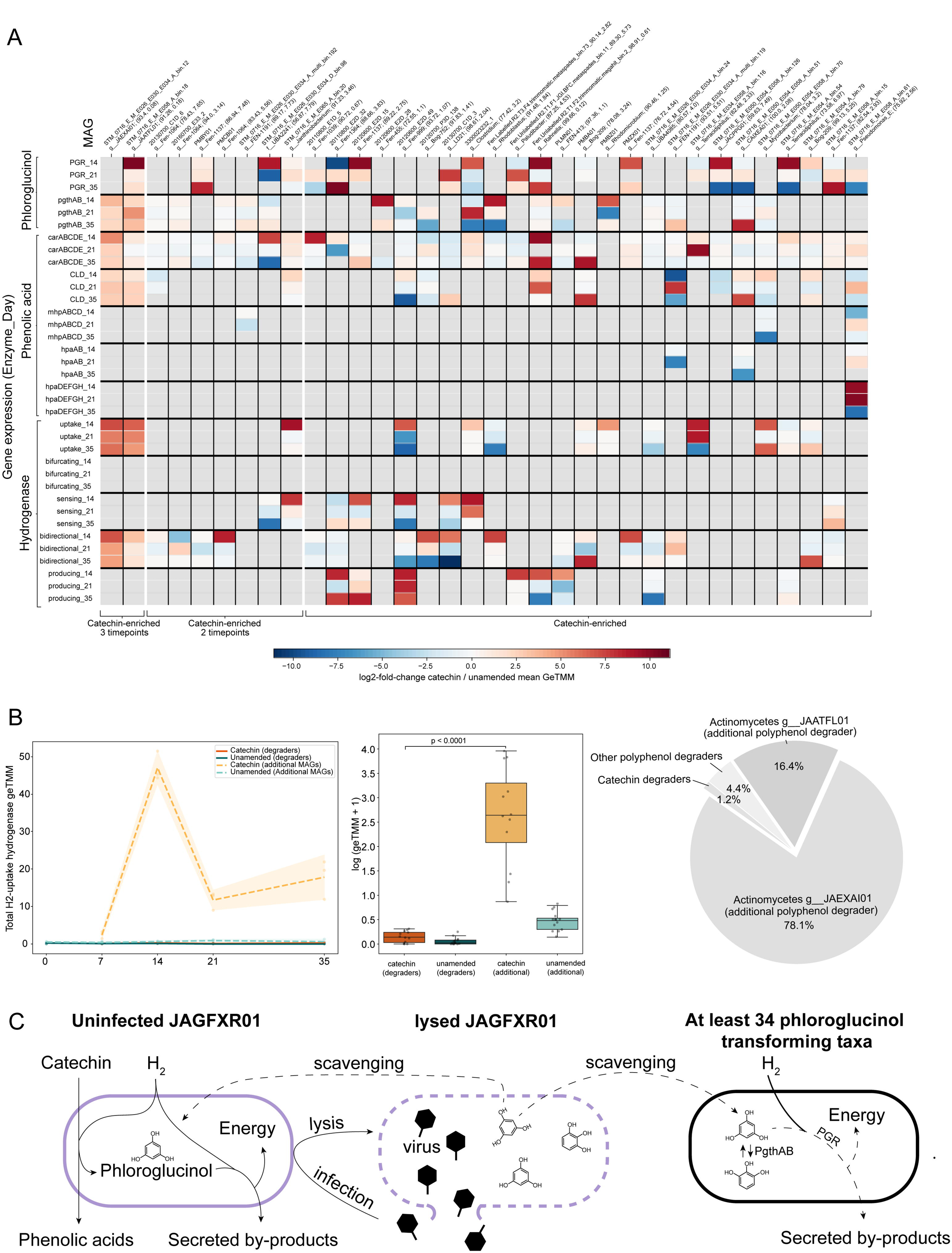
Proposed mechanism by which JAGFXR01-infecting vOTUs facilitate broader microbial community metabolism. **(A)** Heatmap of log2-fold-change expression (GeTMM) between catechin-amended and unamended samples of polyphenol metabolism from additional polyphenol degraders. Rows are gene expression on a specific sample day. Columns are MAGs that did not express genes for catechin-specific degradation. Below each MAG ID is the GTDB genus classification (r214), CheckM2 completeness, and CheckM2 contamination. High quality MAGs with >90% completeness and <5% contamination are bolded. Related metabolisms (phloroglucinol, phenolic acid, or hydrogenase) are grouped together. Hydrogenase gene expression was grouped by direction. Log2-fold-change was computed by adding a pseudocount equal to the minimum magnitude of GeTMM relative abundance (10e-4), taking the mean of sample replicates within a treatment, then dividing the means between treatments that were taken on the same day. The GeTMM values were summed within each sample for genes with multiple subunits (e.g. *pgthAB* is the sum of *pgthA* GeTMM expression and *pgthB* GeTMM expression). Warm colors indicate catechin enrichment and cool colors indicate unamended enrichment. Grey indicates no detection in either catechin-amended or unamended samples. **(B)** Total uptake hydrogenase expression by catechin degraders (solid lines) and additional polyphenol degraders (dashed lines) shown as a time series, boxplot, and pie chart. For the boxplot, a pseudocount of 1 was added to the total geTMM values in each grouping for each sample. The *P* value in the boxplot was derived from a one-sided Welch’s t-test comparing total uptake hydrogenase gene expression between the two MAG groups. **(C)** The proposed mechanism: lysed JAGFXR01 releases partially metabolized catechin intermediates, including compounds that undergo phloroglucinol transformation, enabling their utilization by downstream community members. Solid lines represent interactions supported by the results of this study, and dashed lines represent hypothesized interactions that need to be experimentally evaluated in future studies. JAGFXR01 metabolism is based on previous work [17].

Together, these observations support an expanded model of community carbon metabolism rewiring under catechin amendment by two potential mechanisms. The first proposed expansion is inclusion of a virus shunt mechanism [88, 89], whereby catechin degraders supported the majority of lytic virus activity, leading to phage-mediated lysis and release of bioavailable aromatic catechin-degradation intermediates in cell lysates that stimulated intermediate-specific metabolic activity, and presumably growth, by catechin-degraders and additional polyphenol degraders. The second proposed expansion is that the primary community members that outcompeted methanogens for hydrogen were two relatively rare additional polyphenol degraders (**Figure 5C**). However, these MAGs, too, may have been heavily predated by viruses, as evidenced by the putative phage-host linkage to JAEXAI01 in catechin day 14 samples (**Figure 2C**).

However, alternative hypotheses also could explain phloroglucinol and hydrogen-uptake metabolism enrichment in additional polyphenol degraders. *First*, additional polyphenol-degrading MAGs may have upregulated alternative, non-catechin polyphenol metabolisms that produce or consume pyrogallol and/or phloroglucinol in response to changing redox conditions. To evaluate the first alternative, we examined whether the 34 enriched non-catechin-degrading MAGs encoding *PGR* and/or *pgthAB* also harbored additional polyphenol metabolism genes with differential expression in catechin-amended samples. While 29 of the 34 MAGs showed elevated expression of at least one additional polyphenol-related enzyme during days 14-35 (**Figure S13**), none of these pathways are currently known to be linked to catechin, phloroglucinol, or pyrogallol transformation [90]. *Second*, these inferences are from mostly near-complete MAGs, and so we remain open to the possibility that these MAGs might encode catechin degradation genes that are either novel and so undetected or systematically (for all 34 MAGs to lack the genes) not captured in the partial MAGs. This second alternative hypothesis might also explain why we do not detect upregulated *PGR* expression in additional polyphenol degraders, as they may be utilizing unresolved pathways to metabolize phloroglucinol or pyrogallol. *Third,* host prediction is currently the weakest part of the virus ecogenomics toolkit [57] and while we used only the high-confidence (>90) predictions, false positives for highly expressed phages could alter our conclusions regarding which hosts are heavily predated by lytic phage. However, we did make every effort to increase confidence for some alignment-based predictions (i.e., contig_591846) based on manual inspection of the detailed tool output.

Importantly, our favored proposed explanation of a virus shunt mechanism and the described alternative mechanisms are not mutually exclusive. Regardless of the true mechanism(s), these data-driven mechanistic hypotheses provide clear targets for future experimental testing via (i) long-read sequencing (to complete MAGs), (ii) AI-enabled functional annotation (to improve catechin degradation gene identification), and/or (iii) technically challenging additional experimental measurements such as proximity ligation with appropriate controls [79] (to link viruses to hosts) or stable-isotope probing with ^13^C-catechin that is particularly challenging for viral dynamics [67, 91–93] (to experimentally evaluate which taxa utilize catechin degradation intermediates).

## Conclusion

Overall, these data support an expanded ecosystem-specific model in which catechin amendment of a peat microcosm led to the enrichment of not only microorganisms capable of degrading catechin, but also their viruses. Critically, these viruses were highly transcribed and present in high copy numbers compared to their predicted catechin-degrading hosts, and may be meaningful contributors to microbiome community metabolic rewiring through intense viral predation. Additionally, our results support a model where suppression of the dominant methanogenesis pathway was primarily driven by a narrow subset of two additional polyphenol degraders. These findings extend our previous microbe-focused model of metabolic rewiring [17] to also now recognize a second layer of biological control imposed by viruses. This viral dimension is not simply an overlay on microbial processes; it introduces new causal pathways—through predation, population regulation, and the release of partially transformed metabolites—that might redistribute resources and shuffle the deck on which taxa benefit from catechin amendment. This integrated virus–microbe framework thus offers a hypothesis for how rapid substrate perturbations could drive microbiome community-level metabolic transitions.

More broadly, these findings emphasize the need to consider virus responses as possible unexpected consequences in other ‘perturbed’ ecosystems. For example, our study design, while in soils, mirrors that of diet and nutrition interventions conducted in mammals to tune dysbiotic gut microbiomes back towards healthy states [94, 95]. Given our findings and the well-documented enrichment of prophages in gut microbiota [96, 97], virus induction responses to these interventions will undoubtedly alter microbiome community structure and functioning in ways that need to be understood. We suggest that viruses must be considered beyond their therapeutic potential for targeted microbiome manipulation [97–99] as they are clearly also endemic community members with distinct ecologies that respond dynamically to nutrient enrichments—dynamics that remain poorly understood and underexplored in other systems.

While previous studies have characterized community-level virus responses to perturbation, our approach advances the field by providing a toolkit and conceptual framework to examine the underlying virus-host dynamics that mechanistically drive microbiome functioning and responses.

## Acknowledgements

We are grateful to the EMERGE Biology Integration Institute’s field team for collecting peat samples that were used in the bioreactors and photographs of the site, as well as years of dedicated field support to make the longer-term ecosystem-scale data available. We also thank members of the Sullivan Lab for constructive conversations and suggestions through several years of lab meeting presentations, John Bouranis for early project meeting suggestions, and Simon Roux for troubleshooting iPHoP custom database construction.

## Author contributions

**James Riddell V** (Conceptualization, Data curation, Formal analysis, Investigation, Funding acquisition, Methodology, Project administration, Software, Validation, Visualization, Writing-original draft preparation, Writing- review and editing), **Rokaiya Nurani Shatadru** (Conceptualization, Validation, Methodology, Visualization, Writing — drafting short sections, Writing — review & editing), **Garrett J Smith** (Conceptualization, Validation, Methodology, Writing — review & editing), **Bridget B McGivern** (Conceptualization, Data curation, Methodology, Writing — review & editing), **Jared B Ellenbogen** (Conceptualization, Investigation, Writing — review & editing), **Sophie K Jurgensen** (Conceptualization, Writing — review & editing), **Ami Fofana** (Visualization, Writing — review & editing), **Malak M Tfaily** (Conceptualization, Funding Acquisition, Project administration, Supervision, Writing — review & editing), **Kelly C Wrighton** (Conceptualization, Funding Acquisition, Project administration, Supervision, Writing — drafting short sections, Writing — review & editing), **Matthew B Sullivan** (Conceptualization, Funding Acquisition, Methodology, Project administration, Resources, Supervision, Writing — drafting short sections, Writing — review & editing).

## Supplementary data

Supplementary Figures

Supplementary Data (https://zenodo.org/uploads/18406067)

## Data availability

Metatranscriptomic and metagenomic reads used in this current study are available at NCBI under BioProjectID PRJNA1137330. See **Supplementary Data 1** for specific accession numbers. All code required to reproduce the analyses and visualizations used in this publication are available at https://github.com/jamesriddellv/Grantham_Bioreactor, and are described in the README file. **Supplementary Data 1 through 7** are available in Supplementary Data available at https://zenodo.org/records/18406067 [100]. Assembled contigs and viral contigs identified in this study are available at https://zenodo.org/records/17652154 [33]. Sample metadata and intermediate coverage files used in this study that were not included in the supplementary data are available at: https://zenodo.org/records/17652194 [101].

## Supplemental Figure Legends

**Figure S1. Active virus dataset overview. (A)** Dataset origins of vOTUs from the reference database (left) and vOTUs classified as transcriptionally active (right). **(B)** Distribution of active vOTU estimated quality by CheckV and length. Red line denotes the mean active vOTU length (10.13kb). **(C)** Accumulation curve of active vOTUs across all metatranscriptome sample combinations. The cumulative number of unique, active vOTUs was computed for all possible combinations of samples. For each sample size, the curve represented the mean number of cumulative vOTUs, with the shaded area depicting the range between the minimum and maximum values observed across all permutations. Separate accumulation curves were constructed for the unamended and catechin-amended treatments; for both treatments, samples from Day 0 were included. **(D)** Rank abundance curve of active vOTUs. vOTUs classified as provirus ranked separately. For each vOTU, the orange and turquoise points represent the maximum GeTMM value in each set of samples. vOTUs are ranked by maximum GeTMM value in catechin-amended samples. Heatmap designates which datasets each vOTU was identified in to reveal if these vOTUs were previously identified in previous Stordalen Mire virus studies. **(E)** Violin plot of number of samples each vOTU was active in (left) and number of active vOTUs in each sample (right). Dotted lines represent medians.

**Figure S2. Sensitivity analysis of vOTU activity cutoffs. (A)** Rank abundance of maximum vOTU activity across samples by treatment separated by different activity cutoffs. Cutoffs are nested, so all vOTUs that meet the ≥5 reads mapped to lytic gene also meet the more permissive thresholds. The most conservative threshold is the presented in the main results of this study. **(B-E)** Each row represents a different cutoff used specified on the left side of the figure. Each column is a diversity metric used in this study: **(B)** Species accumulation curve; **(C)** Alpha Diversity; **(D)** Beta Diversity; **(E)** Relative abundance grouped by predicted host.

**Figure S3. Bray-Curtis Dissimilarity**. Sample relative abundance of GeTMM values were normalized with Hellinger transformation. Bray-Curtis dissimilarity between samples were computed for transformed relative abundance values using the vegan package in R. Lighter colors (white, yellow) represent lower dissimilarity (higher similarity) between samples, and darker colors (orange, red) represent higher dissimilarity.

**Figure S4. MAG diversity metrics. (A)** Alpha diversity (Hill order = 1, exp(H’)) of active MAGs in unamended (teal) and catechin-amended (orange) samples. Diversity was computed based on MAG geTMM relative activity values. *P* value based on one-sided Welch’s t-test comparing alpha diversity values z-score standardized within each timepoint between unamended and catechin-amended samples day 7-35. **(B)** Beta diversity represented by PCA of Bray-Curtis dissimilarity between samples based on MAG GeTMM relative activity in each sample. 95% confidence intervals are shown for three distinct sample clusters: catechin and unamended day 7, unamended day 14-35, and catechin day 14-35. These three sample clusters were compared with PERMANOVA (*P* = 0.001). **(C)** Boxplot of Bray-Curtis dissimilarity of sample replicates between treatments. Letters above each boxplot indicate significant differences (Wilcoxon: *P_adj_* < 0.05) in the dissimilarity between days. Days with the same letter within each group (MAG or vOTU) mean the dissimilarity was not statistically different (e.g., dissimilarity between catechin and unamended day 14 samples was not significantly different from the dissimilarity between catechin and unamended day 35 samples).

**Figure S5. Methanogen infection network and relative activity**. (**A**) Methanogen infection network. vOTU-MAG linkages made with iPHoP. Edges represent predictions where the confidence score ≥90. Pink nodes represent active vOTUs/MAGs. (**B**) GeTMM relative activity profiles of vOTUs and MAGs by host genus. vOTU GeTMM values represented on the left axis and MAG GeTMM values represented on the right axis for better comparison. vOTUs are solid lines and MAGs are dashed lines.

**Figure S6. Stacked area chart of active vOTU total reads mapped (left) and GeTMM relative activity (right) in catechin-amended samples summed by predicted host genus.** N specifies how many vOTUs were summed in each group. Aggregated genera with <5% contribution in at least one set of sample replicates were grouped into “Other”.

**Figure S7. Gene-sharing network of highly expressed vOTUs (bold font weight) without a host prediction identified in Fig. 2C and related active vOTUs.** vOTUs are colored by iPHoP-predicted host genus. Grey nodes represent vOTUs with no host assignment by iPHoP. Predicted hosts are labeled whether they were active, inactive, or undetected (iPHoP reference MAG) in microcosm samples. Edges represent family (dashed), subfamily (solid), and genus (double) taxonomic clustering among vOTUs.

**Figure S8. vOTU and MAG relative activity and relative abundance profiles per key carbon cycling genus.** Left y-axis marks vOTU relative abundance values, and right y-axis marks MAG relative abundance values. Solid lines represent vOTUs, dashed lines represent MAGs. vOTU lines are colored by their indicator species analysis cluster combination code from Fig. 2D. For each plot, right to left, top to bottom: unamended metaG, unamended metaT, catechin-amended metaG, catechin-amended metaT **(A)** Clostridium, **(B)** JAGFXR01, **(C)** Paludibacter, **(D)** Pseudomonas_E.

**Figure S9. Heatmap of correlation coefficients between indicator vOTUs and all MAGs in each of four key carbon cycler genera within the peat microcosms.** Black boxes indicate iPHoP-predicted vOTU-MAG pair. Warm colors indicator positive correlation, and cool colors negative correlation. Stars designate statistically significant correlations (BH-adjusted p-value <0.05). **(A)** JAGFXR01, (B) *Clostridium*, (C) *Paludibacter*, (D) *Clostridium*.

**Figure S10. Genomic comparison of vOTUs predicted to infect JAGFXR01 MAGs. (A)** Gene-sharing network of vOTUs predicted to infect JAGFXR01 MAGs using vConTACT3. Nodes represent vOTUs, and edges between nodes represent shared genes. The thickness is the number of genes shared, ranging from 1 to 44, and the size of the node corresponds to the length, ranging from 5.9-59.7Kbp. Colors represent different novel family-level taxonomic predictions based on vConTACT3. contig_860955 failed to cluster and is not shown despite being predicted to infect JAGFXR01. **(B)** Protein similarity heatmap. Values represent the proportion of protein clusters shared between each pair of contigs computed using VirClust.

**Figure S11. Cumulative relative expression of phloroglucinol reductase (*PGR*) and pyrogallol transhydroxylase (*pgthAB*) in non-catechin degraders (n=104) and catechin degraders.** Total relative expression calculated by summing relative expression across MAGs in each sample for each group of MAGs (catechin vs non-catechin degraders). Number of MAGs in each group indicated in y-axis legends. *P* value significance of Welch’s t-test between catechin and unamended samples day 7-35 shown in the title of each subplot. ns = not significant; * = 0.05; ** = 0.01; *** = 0.001.

**Figure S12. (A)** Total hydrogenase geTMM relative abundance between treatments over time in additional polyphenol degraders (n=34). Hydrogenase gene expression grouped by direction. Total geTMM relative abundance of genes annotated with polyphenol degradation pathways for **(B)** the undescribed *Actinomycetota* g JAEXAI01 and **(C)** the undescribed *Actinomycetota* g JAATFL01. *P* value significance of Welch’s t-test between catechin and unamended samples day 7-35 shown in the title of each subplot. ns = not significant; * = 0.05; ** = 0.01; *** = 0.001.

