## Supplemental Figures for "Viruses help shape microbiome response to polyphenol rewiring of methane-suppressed peat microcosms"

### Supplementary Figures

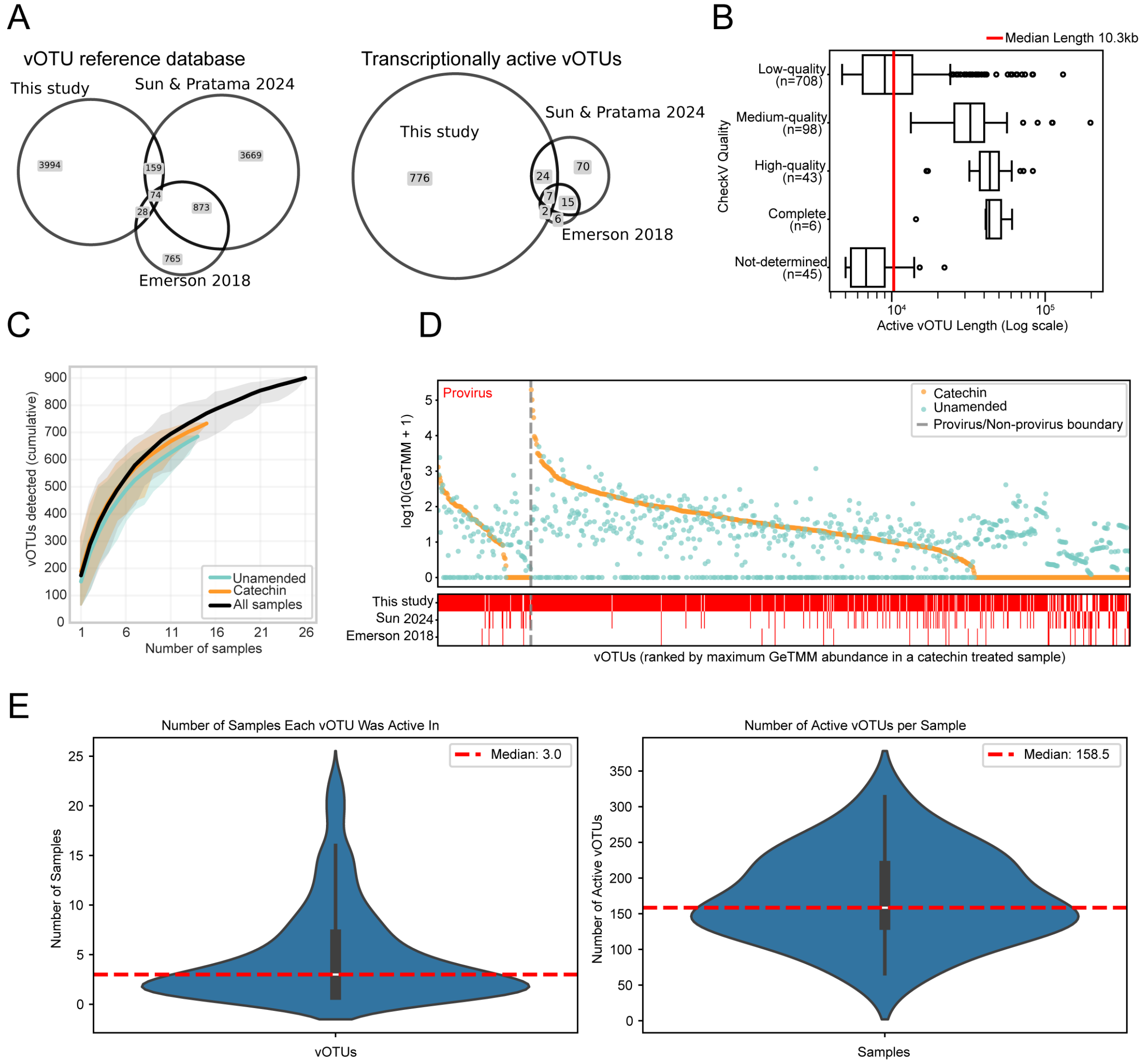


**Figure S1. Active virus dataset overview. (A)** Dataset origins of vOTUs from the reference database (left) and vOTUs classified as transcriptionally active (right). **(B)** Distribution of active vOTU estimated quality by CheckV and length. Red line denotes the mean active vOTU length (10.13kb). **(C)** Accumulation curve of active vOTUs across all metatranscriptome sample combinations. The cumulative number of unique, active vOTUs was computed for all possible combinations of samples. For each sample size, the curve represented the mean number of cumulative vOTUs, with the shaded area depicting the range between the minimum and maximum values observed across all permutations. Separate accumulation curves were constructed for the unamended and catechin-amended treatments; for both treatments, samples from Day 0 were included. **(D)** Rank abundance curve of active vOTUs. vOTUs classified as provirus ranked separately. For each vOTU, the orange and turquoise points represent the maximum GeTMM value in each set of samples. vOTUs are ranked by maximum GeTMM value in catechin-amended samples. Heatmap designates which datasets each vOTU was identified in to reveal if these vOTUs were previously identified in previous Stordalen Mire virus studies. **(E)** Violin plot of number of samples each vOTU was active in (left) and number of active vOTUs in each sample (right). Dotted lines represent medians.

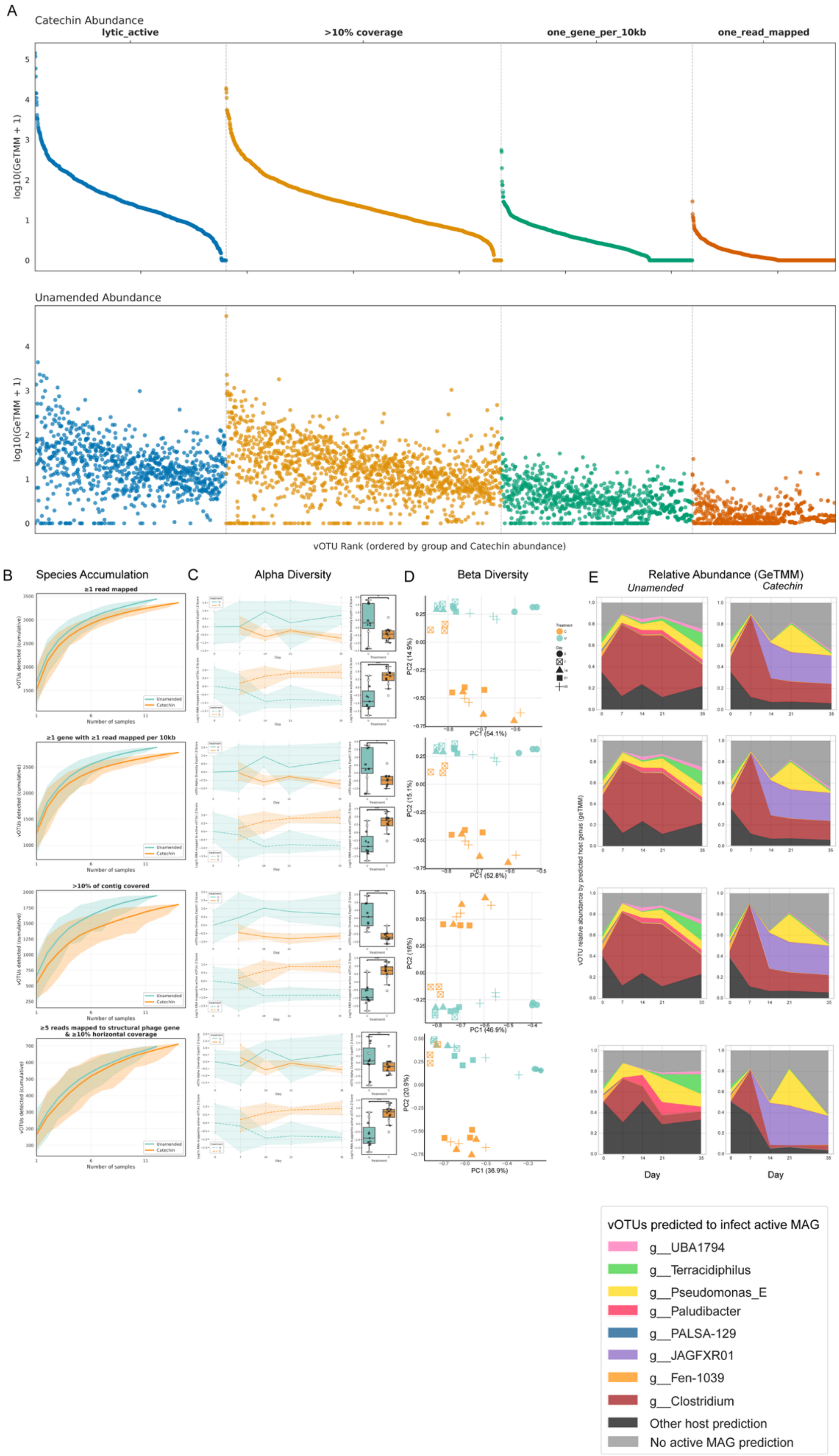
A
Catechin Abundance
lytic_active
>10% coverage
one_gene_per_10kb
one_read_mapped
log10(GeTMM + 1)
Unamended Abundance
log10(GeTMM + 1)
vOTU Rank (ordered by group and Catechin abundance)
B
Species Accumulation
≥1 read mapped
≥1 gene with ≥1 read mapped per 10kb
>10% of contig covered
≥5 reads mapped to structural phage gene & ≥10% horizontal coverage
Number of samples
Unamended
Catechin
C
Alpha Diversity
D
Beta Diversity
PC1 (54.1%)
PC2 (14.9%)
PC1 (52.8%)
PC2 (15.1%)
PC1 (46.9%)
PC2 (16%)
PC1 (36.9%)
PC2 (20.9%)
E
Relative Abundance (GeTMM)
Unamended
Catechin
vOTU relative abundance by predicted host genus (geTMM)
Day
Day
vOTUs predicted to infect active MAG
g__UBA1794
g__Terracidiphilus
g__Pseudomonas_E
g__Paludibacter
g__PALSA-129
g__JAGFXR01
g__Fen-1039
g__Clostridium
Other host prediction
No active MAG prediction

**Figure S2. Sensitivity analysis of vOTU activity cutoffs. (A)** Rank abundance of maximum vOTU activity across samples by treatment separated by different activity cutoffs. Cutoffs are nested, so all vOTUs that meet the ≥5 reads mapped to lytic gene also meet the more permissive thresholds. The most conservative threshold is the presented in the main results of this study. **(B-E)** Each row represents a different cutoff used specified on the left side of the figure. Each column is a diversity metric used in this study: **(B)** Species accumulation curve; **(C)** Alpha Diversity; **(D)** Beta Diversity; **(E)** Relative abundance grouped by predicted host.

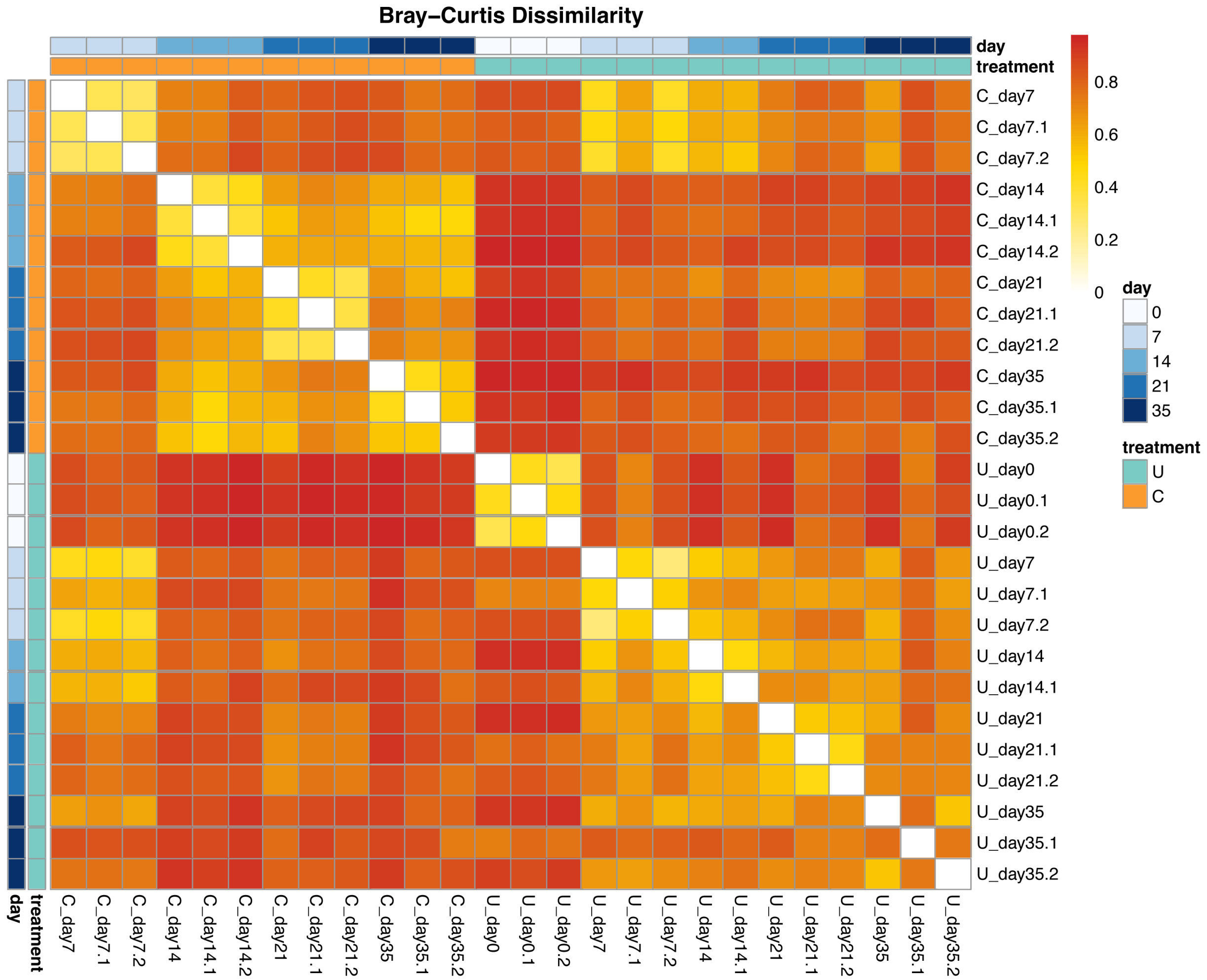


**Figure S3. Bray-Curtis Dissimilarity**. Sample relative abundance of GeTMM values were normalized with Hellinger transformation. Bray-Curtis dissimilarity between samples were computed for transformed relative abundance values using the vegan package in R. Lighter colors (white, yellow) represent lower dissimilarity (higher similarity) between samples, and darker colors (orange, red) represent higher dissimilarity.

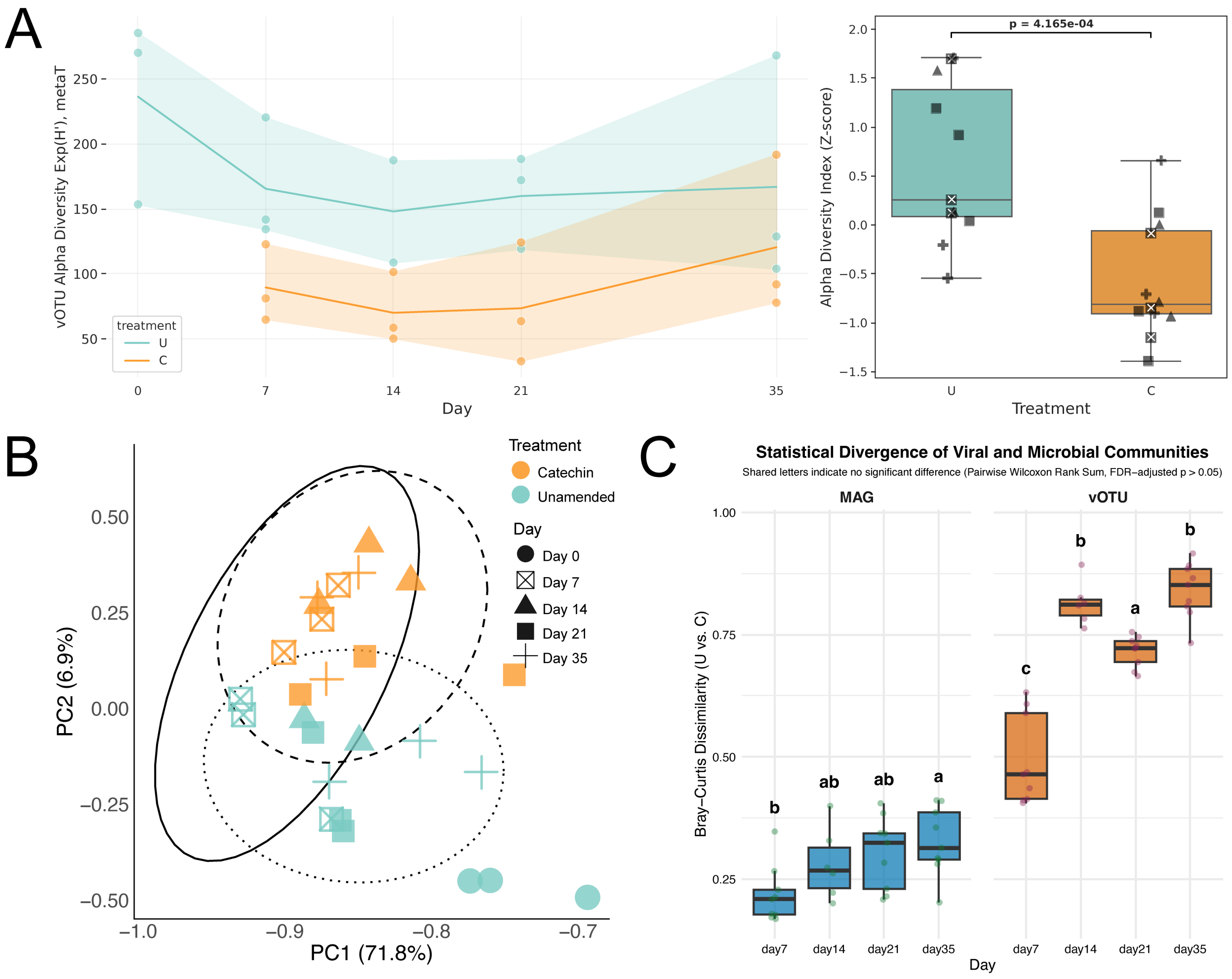


**Figure S4. MAG diversity metrics. (A)** Alpha diversity (Hill order = 1, exp(H')) of active MAGs in unamended (teal) and catechin-amended (orange) samples. Diversity was computed based on MAG geTMM relative activity values. *P* value based on one-sided Welch's t-test comparing alpha diversity values z-score standardized within each timepoint between unamended and catechin-amended samples day 7-35. **(B)** Beta diversity represented by PCA of Bray-Curtis dissimilarity between samples based on MAG GeTMM relative activity in each sample. 95% confidence intervals are shown for three distinct sample clusters: catechin and unamended day 7, unamended day 14-35, and catechin day 14-35. These three sample clusters were compared with PERMANOVA (*P* = 0.001). **(C)** Boxplot of Bray-Curtis dissimilarity of sample replicates between treatments. Letters above each boxplot indicate significant differences (Wilcoxon: $P_{adj} < 0.05$) in the dissimilarity between days. Days with the same letter within each group (MAG or vOTU) mean the dissimilarity was not statistically different (e.g., dissimilarity between catechin and unamended day 14 samples was not significantly different from the dissimilarity between catechin and unamended day 35 samples).

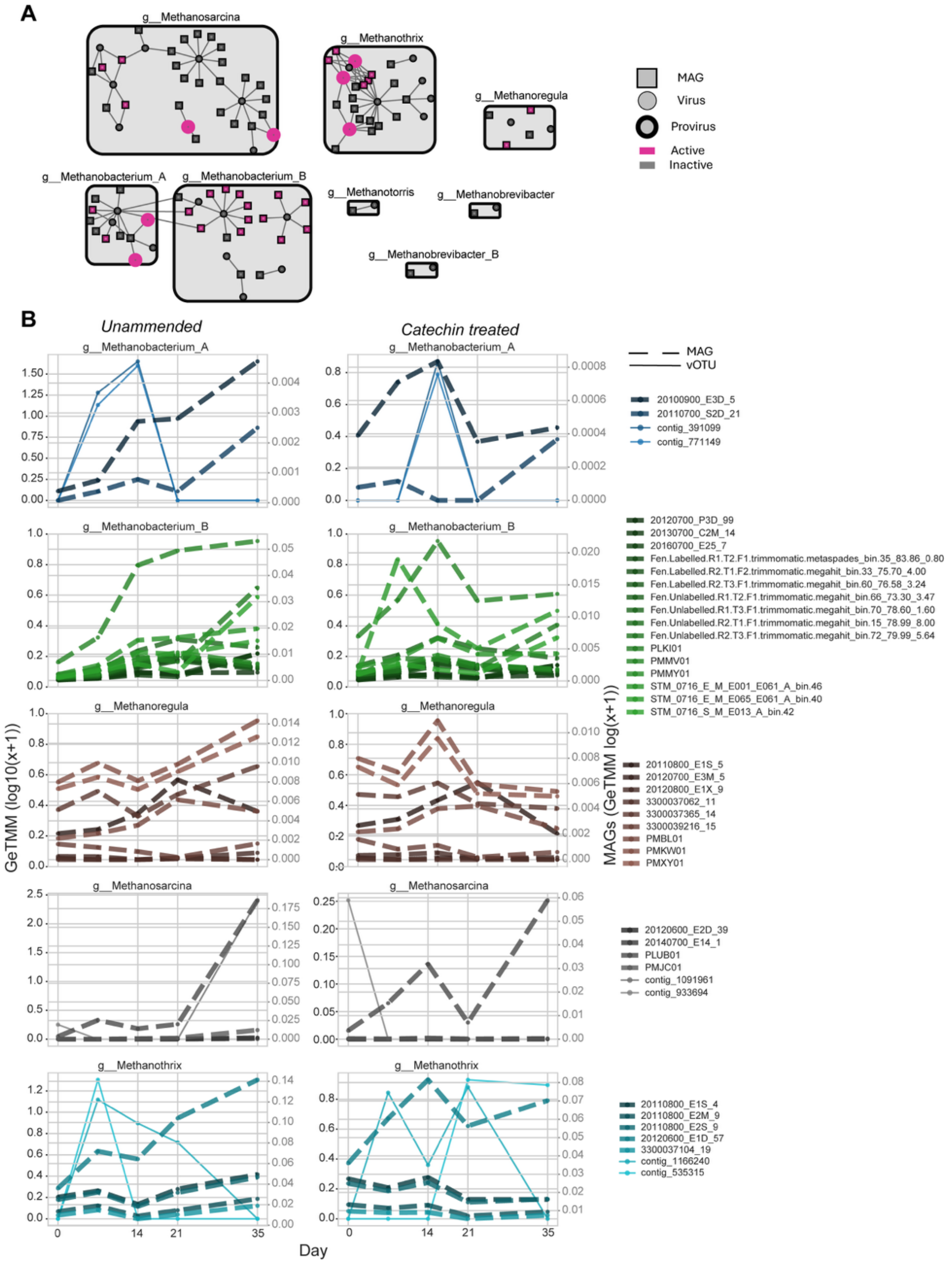


**Figure S5. Methanogen infection network and relative activity**. (**A**) Methanogen infection network. vOTU-MAG linkages made with iPHoP. Edges represent predictions where the confidence score ≥90. Pink nodes represent active vOTUs/MAGs. (**B**) GeTMM relative activity profiles of vOTUs and MAGs by host genus. vOTU GeTMM values represented on the left axis and MAG GeTMM values represented on the right axis for better comparison. vOTUs are solid lines and MAGs are dashed lines.

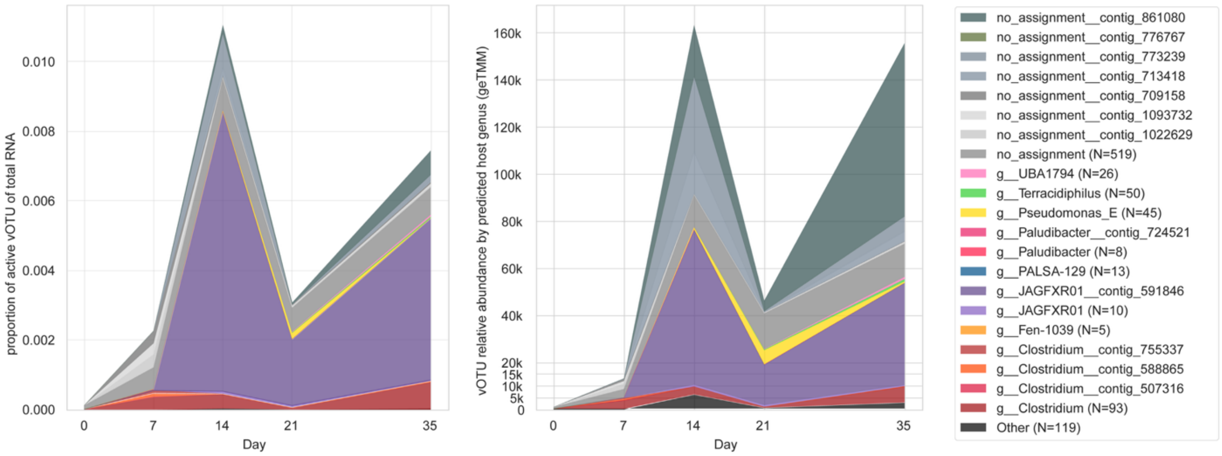


**Figure S6. Stacked area chart of active vOTU total reads mapped (left) and GeTMM relative activity (right) in catechin-amended samples summed by predicted host genus.** N specifies how many vOTUs were summed in each group. Aggregated genera with <5% contribution in at least one set of sample replicates were grouped into “Other”.

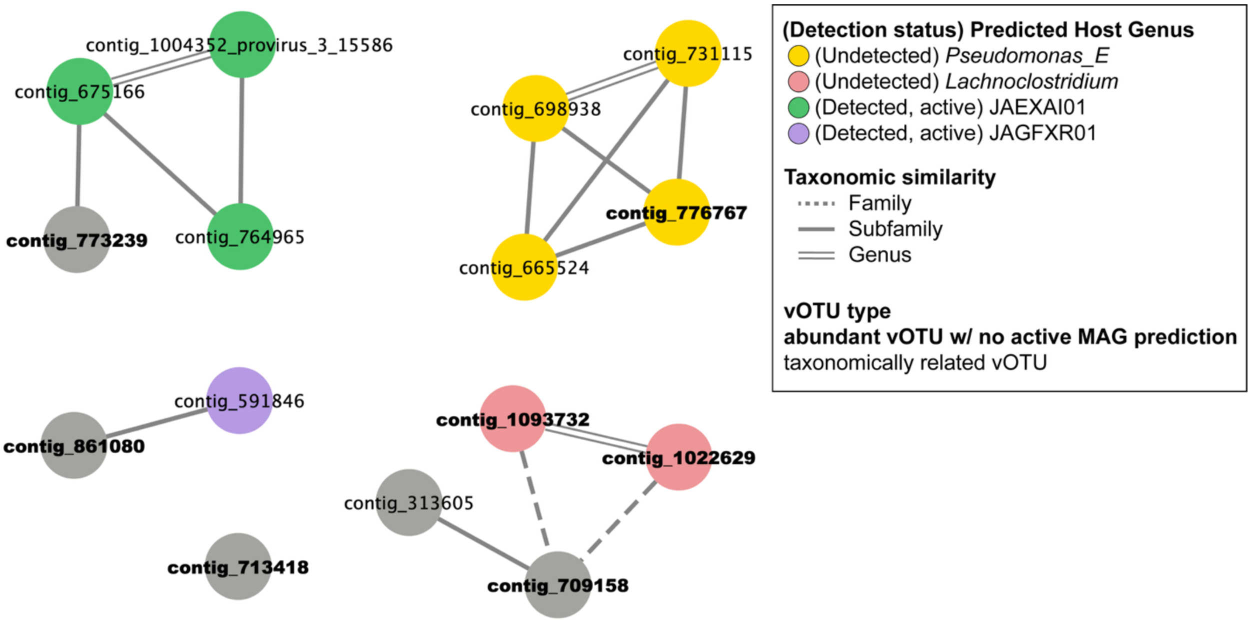


**Figure S7. Gene-sharing network of highly expressed vOTUs (bold font weight) without a host prediction identified in Fig. 2C and related active vOTUs.** vOTUs are colored by iPHoP-predicted host genus. Grey nodes represent vOTUs with no host assignment by iPHoP. Predicted hosts are labeled whether they were active, inactive, or undetected (iPHoP reference MAG) in microcosm samples. Edges represent family (dashed), subfamily (solid), and genus (double) taxonomic clustering among vOTUs.

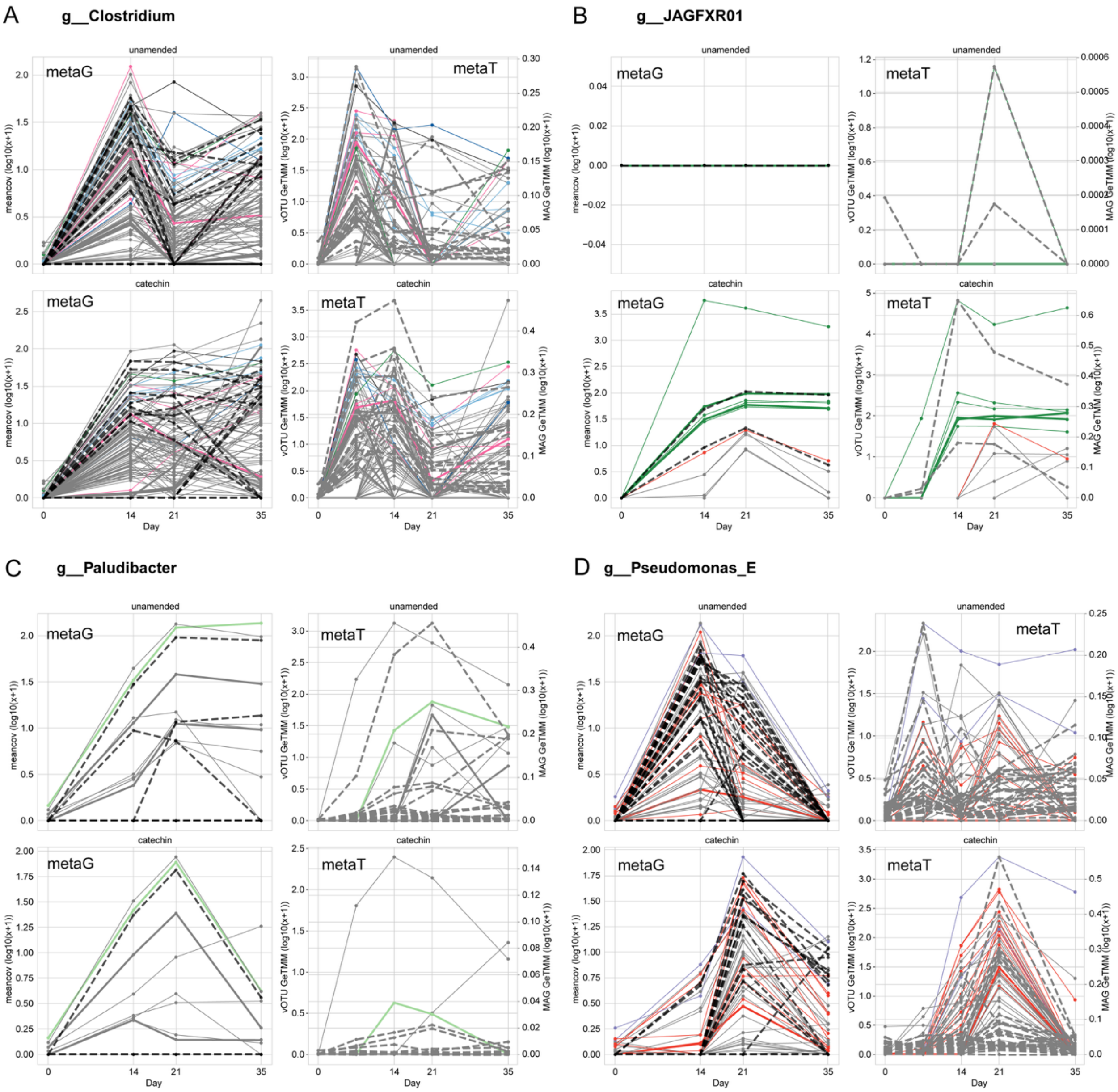


**Figure S8. vOTU and MAG relative activity and relative abundance profiles per key carbon cycling genus.** Left y-axis marks vOTU relative abundance values, and right y-axis marks MAG relative abundance values. Solid lines represent vOTUs, dashed lines represent MAGs. vOTU lines are colored by their indicator species analysis cluster combination code from Fig. 2D. For each plot, right to left, top to bottom: unamended metaG, unamended metaT, catechin-amended metaG, catechin-amended metaT **(A)** Clostridium, **(B)** JAGFXR01, **(C)** Paludibacter, **(D)** Pseudomonas_E.

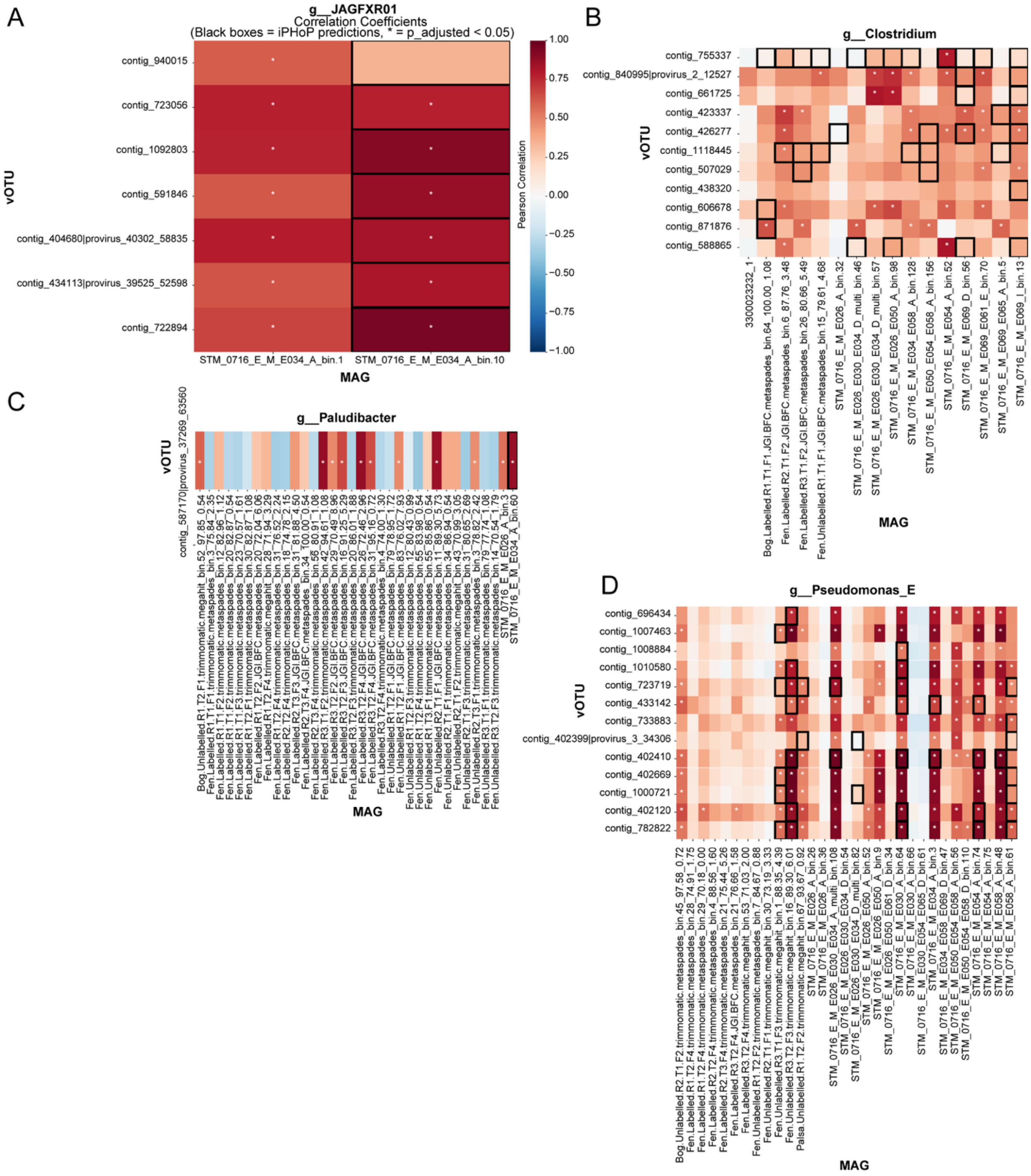


**Figure S9. Heatmap of correlation coefficients between indicator vOTUs and all MAGs in each of four key carbon cycler genera within the peat microcosms.** Black boxes indicate iPHoP-predicted vOTU-MAG pair. Warm colors indicator positive correlation, and cool colors negative correlation. Stars designate statistically significant correlations (BH-adjusted p-value <0.05). **(A)** JAGFXR01, **(B)** *Clostridium*, **(C)** *Paludibacter*, **(D)** *Clostridium*.

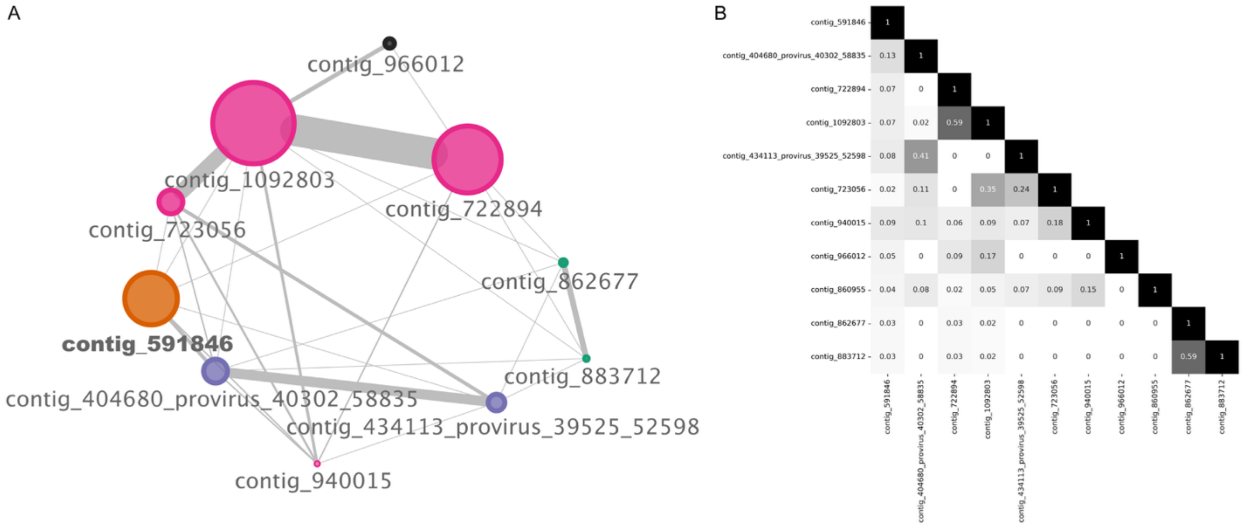


**Figure S10. Genomic comparison of vOTUs predicted to infect JAGFXR01 MAGs. (A)** Gene-sharing network of vOTUs predicted to infect JAGFXR01 MAGs using vConTACT3. Nodes represent vOTUs, and edges between nodes represent shared genes. The thickness is the number of genes shared, ranging from 1 to 44, and the size of the node corresponds to the length, ranging from 5.9-59.7Kbp. Colors represent different novel family-level taxonomic predictions based on vConTACT3. contig_860955 failed to cluster and is not shown despite being predicted to infect JAGFXR01. **(B)** Protein similarity heatmap. Values represent the proportion of protein clusters shared between each pair of contigs computed using VirClust.

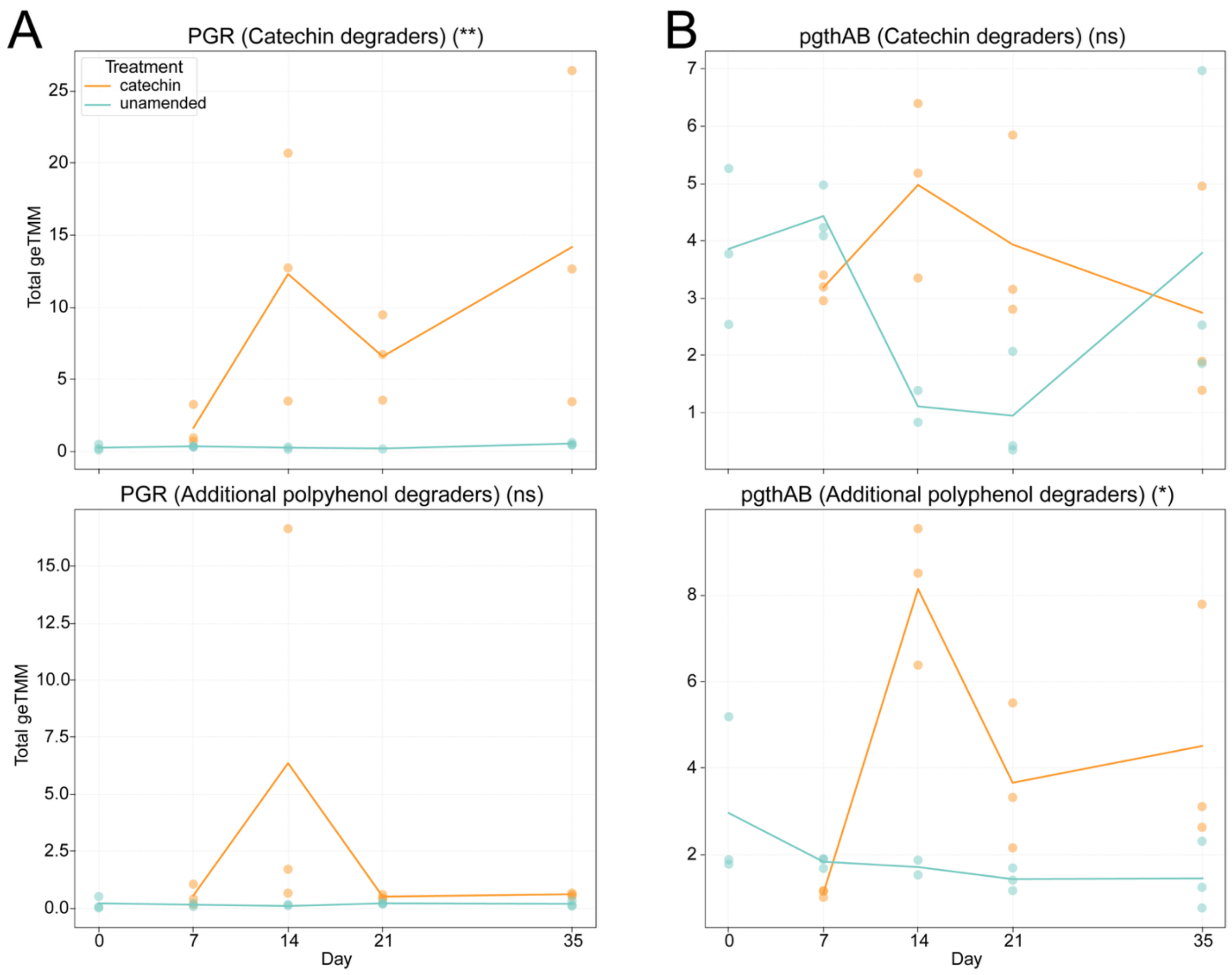


**Figure S11. Cumulative relative expression of phloroglucinol reductase (*PGR*) and pyrogallol transhydroxylase (*pgthAB*) in non-catechin degraders (n=104) and catechin degraders.** Total relative expression calculated by summing relative expression across MAGs in each sample for each group of MAGs (catechin vs non-catechin degraders). Number of MAGs in each group indicated in y-axis legends. *P* value significance of Welch's t-test between catechin and unamended samples day 7-35 shown in the title of each subplot. ns = not significant; * = 0.05; ** = 0.01; *** = 0.001.

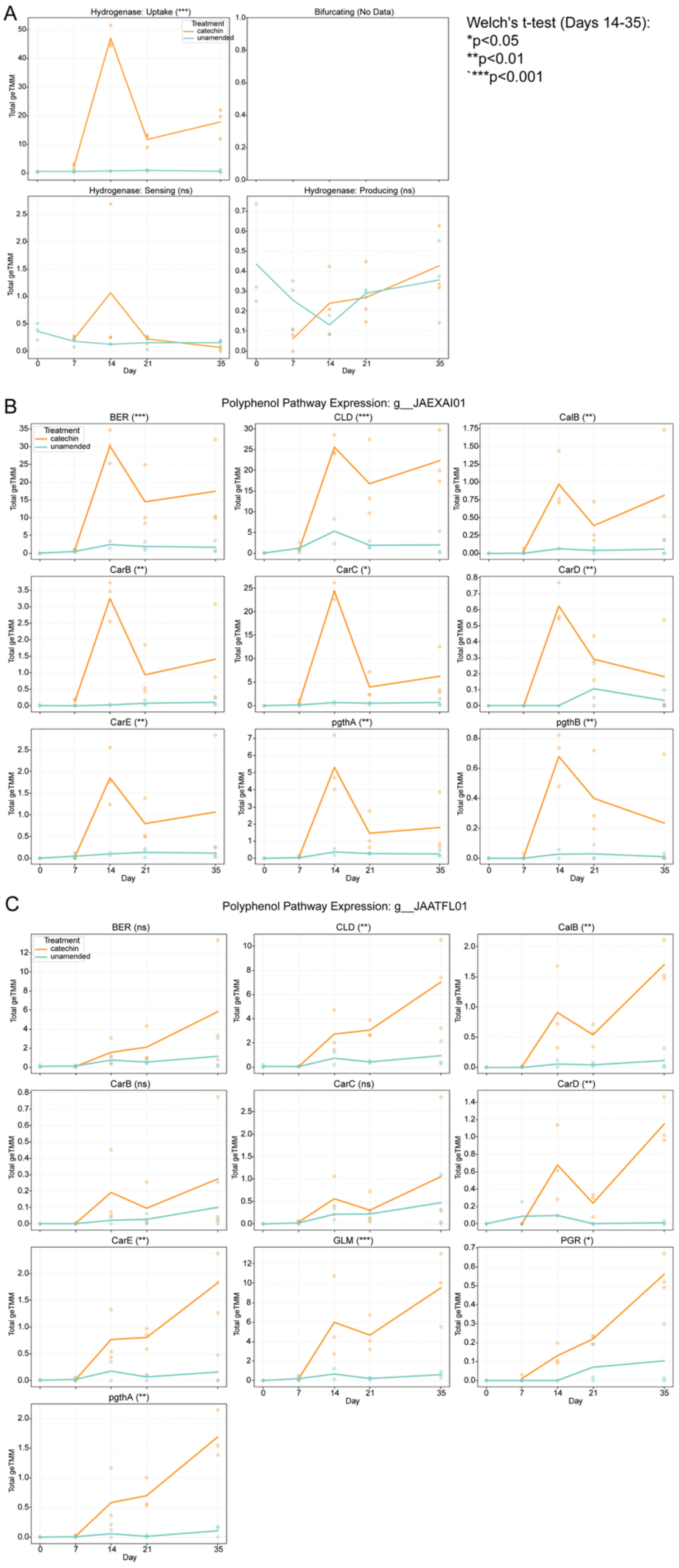
A
Hydrogenase: Uptake (***)
Bifurcating (No Data)
Hydrogenase: Sensing (ns)
Hydrogenase: Producing (ns)
Treatment
catechin
unamended
Total geTMM
Day
Welch's t-test (Days 14-35):
*p<0.05
**p<0.01
`***p<0.001
B
Polyphenol Pathway Expression: g__JAEXAI01
BER (***)
CLD (***)
CalB (**)
CarB (**)
CarC (*)
CarD (**)
CarE (**)
pgthA (**)
pgthB (**)
C
Polyphenol Pathway Expression: g__JAATFL01
BER (ns)
CLD (**)
CalB (**)
CarB (ns)
CarC (ns)
CarD (**)
CarE (**)
GLM (***)
PGR (*)
pgthA (**)

**Figure S12. (A)** Total hydrogenase geTMM relative abundance between treatments over time in additional polyphenol degraders (n=34). Hydrogenase gene expression grouped by direction. Total geTMM relative abundance of genes annotated with polyphenol degradation pathways for **(B)** the undescribed *Actinomycetota* g__JAEXAI01 and **(C)** the undescribed *Actinomycetota* g__JAATFL01. *P* value significance of Welch's t-test between catechin and unamended samples day 7-35 shown in the title of each subplot. ns = not significant; * = 0.05; ** = 0.01; *** = 0.001.

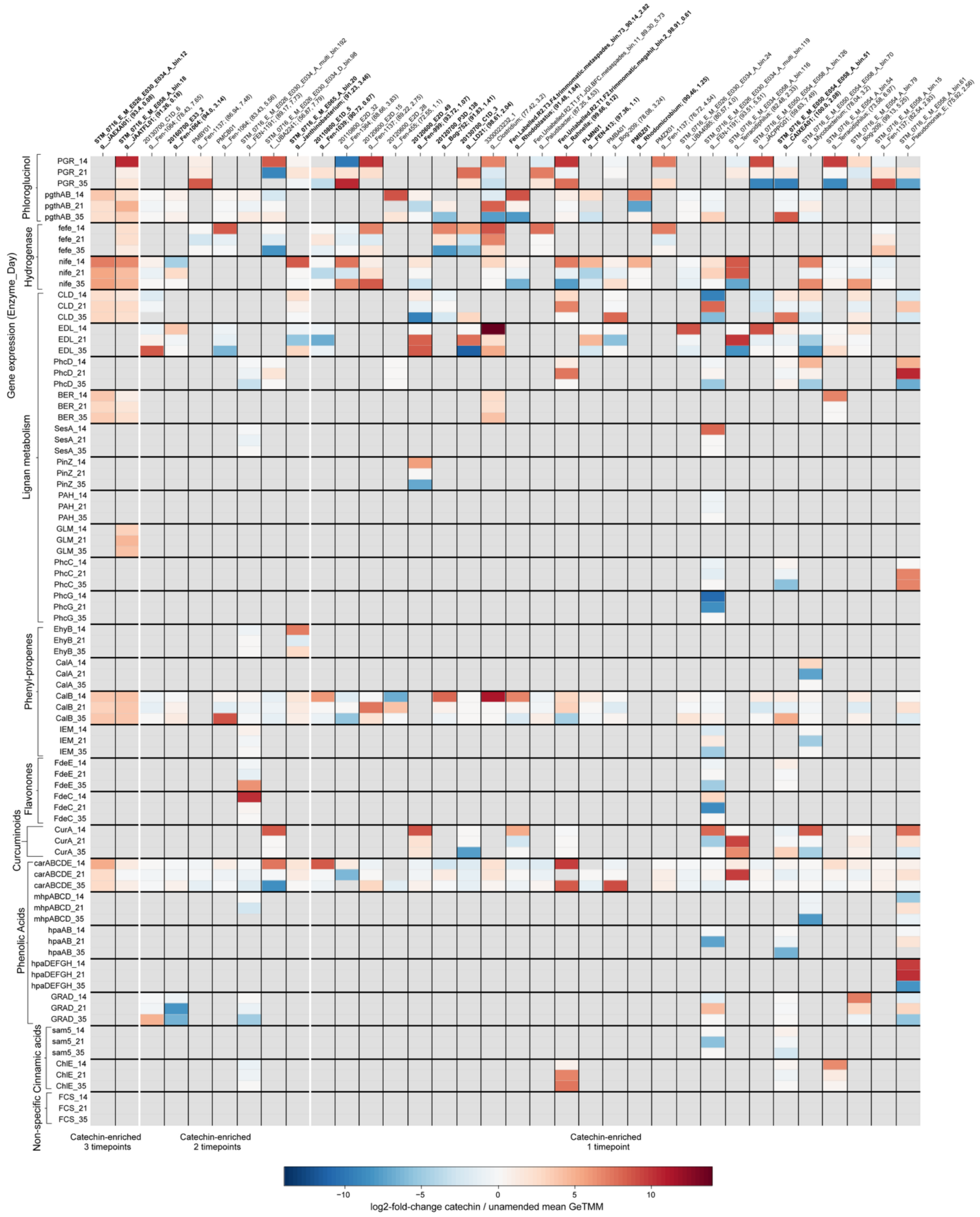


**Figure S13. Extended heatmap of log2-fold-change expression (GeTMM) between catechin-amended and unamended samples of polyphenol metabolism from non-catechin-degraders.** Rows are gene expression on a specific sample day. Columns are MAGs that did not express genes for catechin degradation. Below each MAG ID is the GTDB genus classification (r214), CheckM2 completeness, and CheckM2 contamination. Related metabolisms are grouped together based on the CAMPER metabolism map (https://github.com/WrightonLabCSU/CAMPER/blob/main/figs/camper_map.png).
